# Lagging strand gap suppression connects BRCA-mediated fork protection to nucleosome assembly by ensuring PCNA-dependent CAF-1 recycling

**DOI:** 10.1101/2021.11.08.467732

**Authors:** Tanay Thakar, Joshua Straka, Claudia M. Nicolae, George-Lucian Moldovan

**Affiliations:** Department of Biochemistry and Molecular Biology, The Pennsylvania State University College of Medicine, Hershey, PA 17033, USA

## Abstract

The inability to protect stalled replication forks from nucleolytic degradation drives genome instability and is associated with chemosensitivity in BRCA-deficient tumors. An emerging hallmark of BRCA deficiency is the inability to suppress replication-associated single-stranded DNA (ssDNA) gaps. Here, we report that ssDNA gaps on the lagging strand interfere with the ASF1-CAF-1 pathway of nucleosome assembly, and drive fork degradation in BRCA-deficient cells. We show that CAF-1 function at replication forks is lost in BRCA-deficient cells, due to its sequestration at inactive replication factories during replication stress. This CAF-1 recycling defect is caused by the accumulation of Polα-dependent lagging strand gaps, which preclude PCNA unloading, causing sequestration of PCNA-CAF-1 complexes on chromatin. Importantly, correcting PCNA unloading defects in BRCA-deficient cells restores fork stability in a CAF-1-dependent manner. We also show that the activation of a HIRA-dependent compensatory histone deposition pathway restores fork stability to BRCA-deficient cells upon CAF-1 loss. We thus define nucleosome assembly as a critical determinant of BRCA-mediated fork stability. We further reveal lagging strand ssDNA gaps as drivers of fork degradation in BRCA-deficient cells, which operate by inhibiting PCNA unloading and CAF-1-dependent nucleosome assembly.

## INTRODUCTION

The breast cancer susceptibility factors BRCA1 and BRCA2 act as tumor suppressors, by promoting accurate DNA repair through homologous recombination (HR) and protecting against genomic instability, an enabling hallmark of cancer (Hanahan and Weinberg, 2011; Jasin, 2002). Germline mutations in the BRCA1 and BRCA2 genes drastically increase the lifetime risk of developing breast and ovarian cancer (Kuchenbaecker et al., 2017; Scully and Livingston, 2000; Welcsh et al., 2000). In addition to mediating DNA double strand break (DSB) repair by HR, the BRCA proteins also play a critical role in maintaining the integrity of DNA replication forks during replication stress (Schlacher et al., 2011, 2012). A global response to replication stress is replication fork reversal, which involves the regression of replication forks and the annealing of complementary nascent DNA strands (Neelsen and Lopes, 2015; Quinet et al., 2017a; Zellweger et al., 2015). Fork reversal necessitates the engagement of the BRCA pathway to stabilize nascent DNA via the formation of RAD51 nucleofilaments. In the absence of an intact BRCA pathway, nascent DNA at reversed forks becomes susceptible to degradation by nucleases, namely MRE11, EXO1 and DNA2 (Lemaçon et al., 2017; Ray Chaudhuri et al., 2016; Schlacher et al., 2011, 2012). A direct consequence of fork degradation is the accumulation of DNA damage and gross chromosomal aberrations, making fork protection a major mechanism by which the BRCA pathway maintains genome stability and tumor suppression (Schlacher et al., 2011, 2012). Importantly, fork degradation is also linked to sensitivity to chemotherapeutic agents, and restoration of fork stability is associated with chemoresistance in BRCA deficient cancers (Ray Chaudhuri et al., 2016).

The sliding DNA clamp <u>P</u>roliferating <u>C</u>ell <u>N</u>uclear <u>A</u>ntigen (PCNA) is a core component of the DNA replication machinery. During DNA synthesis, PCNA interacts with DNA polymerases to maintain their engagement on template DNA, thereby increasing their processivity. In addition to polymerase recruitment, PCNA also serves as a scaffold for the recruitment of numerous other replication factors and acts as a functional toolbelt for DNA replication (Choe and Moldovan, 2017). PCNA also orchestrates replication coupled nucleosome assembly by recruiting the histone chaperone <u>C</u>hromatin <u>A</u>ssembly <u>F</u>actor-1 (CAF-1) (Shibahara and Stillman, 1999; Zhang et al., 2000). PCNA performs distinct functions during leading and lagging strand DNA replication: On the leading strand, PCNA supports continuous DNA synthesis by recruiting and facilitating the function of Polε. On the lagging strand, PCNA recruits Polδ to synthesize Okazaki fragments (OFs) by extending RNA primers assembled by Polα. Subsequently, PCNA recruits the flap endonuclease FEN1 to cleave downstream RNA primers displaced by Polδ, and the DNA ligase LIG1 to seal the resulting nick to yield intact stretches of DNA (Balakrishnan and Bambara, 2013).

Precise DNA replication requires a tight regulation of PCNA cycling at replication forks. PCNA is loaded during replication initiation by the RFC1-5 complex and unloaded by an RFC-like complex composed of ATAD5 and RFC2-5 upon replication completion (Kang et al., 2019a; Lee et al., 2013; Majka and Burgers, 2004). On lagging strands, frequent Polα mediated repriming necessitates the constant loading of PCNA homotrimers to support the synthesis of multiple OFs. PCNA is unloaded by ATAD5 from the lagging strand upon OF maturation. OF ligation is an essential prerequisite for PCNA unloading, and a failure to unload PCNA can drive genome instability by sequestering PCNA interacting factors at inactive replication factories (Johnson et al., 2016; Kubota et al., 2015; Lee et al., 2013).

RAD18-mediated ubiquitination of PCNA at the lysine 164 (K164) residue is a prominent response of eukaryotic cells to replication stress. This modification enables the post replicative repair (PRR) of ssDNA gaps through translesion synthesis (TLS) (Hoege et al., 2002; Kannouche et al., 2004; Karras and Jentsch, 2010; Lopes et al., 2006; Stelter and Ulrich, 2003). By generating PCNA-K164R mutant human cell lines, completely deficient in PCNA ubiquitination, we recently uncovered an essential role of ubiquitinated PCNA in preventing the nucleolytic degradation of stalled replication forks (Thakar et al., 2020). Mechanistically, we showed that fork degradation in PCNA-K164R cells is caused by the accumulation of lagging strand gaps, which sequester PCNA as OF ligation is impaired. Since CAF-1 forms a tight complex with PCNA, it is also sequestered in these PCNA complexes in the wake of replication forks, thus impeding replication-coupled nucleosome establishment in these cells, and priming stressed forks for nucleolytic degradation.

Recent publications have revealed a previously underappreciated role of the BRCA-RAD51 pathway in suppressing the accumulation of replication-associated single stranded DNA (ssDNA) gaps. Gap mitigation by the BRCA pathway occurs through two distinct mechanisms: 1) by restraining fork progression during replication stress, thereby suppressing excessive PRIMPOL-mediated fork repriming (Kang et al., 2021; Liu et al., 2020; Panzarino et al., 2021; Simoneau et al., 2021; Taglialatela et al., 2021); and 2) by promoting RAD51-dependent PRR of gaps (Hashimoto et al., 2010; Kolinjivadi et al., 2017; Piberger et al., 2020; Thakar et al., 2020; Tirman et al., 2021). Importantly, replication-associated gaps have been connected to PARP inhibitor (PARPi) sensitivity in BRCA-deficient cells (Cong et al., 2021; Paes Dias et al., 2021; Panzarino et al., 2021; Simoneau et al., 2021; Thakar et al., 2020). Interestingly, Okazaki fragment processing defects have also been identified in BRCA-deficient cells (Cong et al., 2021; Paes Dias et al., 2021).

Here, we show that nucleosome establishment controls fork protection, genomic stability, and chemoresistance of BRCA-deficient cells. Mechanistically, we show that Polα-dependent lagging strand ssDNA gaps cause CAF-1 recycling defects in BRCA-deficient cells, since they sequester PCNA-CAF-1 complexes behind replication forks. The subsequent reduction in CAF-1 availability at ongoing replication forks underlies fork degradation in BRCA-deficient cells. Indeed, we demonstrate that correcting PCNA unloading defects restores fork protection to BRCA-deficient cells in a CAF-1-dependent manner, thereby exposing efficient PCNA unloading and CAF-1-mediated nucleosome establishment as major effectors of fork protection by the BRCA-RAD51 pathway. Moreover, we show that loss of CAF-1 restores fork protection to BRCA-deficient cells by releasing an alternative nucleosome assembly pathway mediated by the histone chaperone HIRA. Our work uncovers an unexpected role for nucleosome deposition pathways in mediating BRCA-dependent genome stability.

## RESULTS

### CAF-1 inactivation rescues fork stability in PCNA-K164R and BRCA-deficient cells

We recently showed that accumulation of replication-associated ssDNA gaps drives fork degradation at stalled replication forks in PCNA-K164R cells, by eliciting defective nucleosome assembly by CAF-1 at replication forks (Thakar et al., 2020). Surprisingly, while knockdown of CHAF1A (the large subunit of the heterotrimeric CAF-1 complex) elicited fork degradation in 293T-WT cells as we previously documented, fork stability was completely rescued in 293T-K164R cells upon CHAF1A depletion (Supplementary Fig. S1A,B). Recent publications have characterized a previously underappreciated role of the BRCA1 and BRCA2 proteins in suppressing replication-coupled ssDNA gaps (Cong et al., 2021; Kolinjivadi et al., 2017; Panzarino et al., 2021; Taglialatela et al., 2021; Thakar et al., 2020). Given that we showed PCNA-ubiquitination and the BRCA-pathway operate in parallel to suppress ssDNA gaps during DNA replication (Thakar et al., 2020), we sought to investigate the effect of CHAF1A inactivation on fork stability in cells deficient in either BRCA1 or BRCA2 function. Strikingly, similarly to PCNA-K164R cells, CHAF1A depletion fully restored fork stability to HeLa-BRCA2^KO^ as well as RPE1-p53^KO^BRCA1^KO^ cells, while causing fork degradation in their respective BRCA-proficient counterparts (Fig. 1A,B; Supplementary Fig. S1C,D). To rule out potential siRNA off-target effects, we employed CRISPR/Cas9 to knock-out CHAF1A in 293T and HeLa cells (Supplementary Fig. S1E,F). Similar to CHAF1A depletion using siRNA, both 293T and HeLa CHAF1A-knockout cells displayed fork degradation upon HU treatment (Fig. 1C,D). Importantly, BRCA2 depletion caused fork degradation in wildtype, but not in CHAF1A-knockout HeLa and 293T cells (Fig. 1C,D; Supplementary Fig. S1G,H). These findings indicate that loss of CAF-1 promotes fork degradation in wildtype cells, but suppresses this degradation in BRCA-deficient cells.

**Figure 1.**
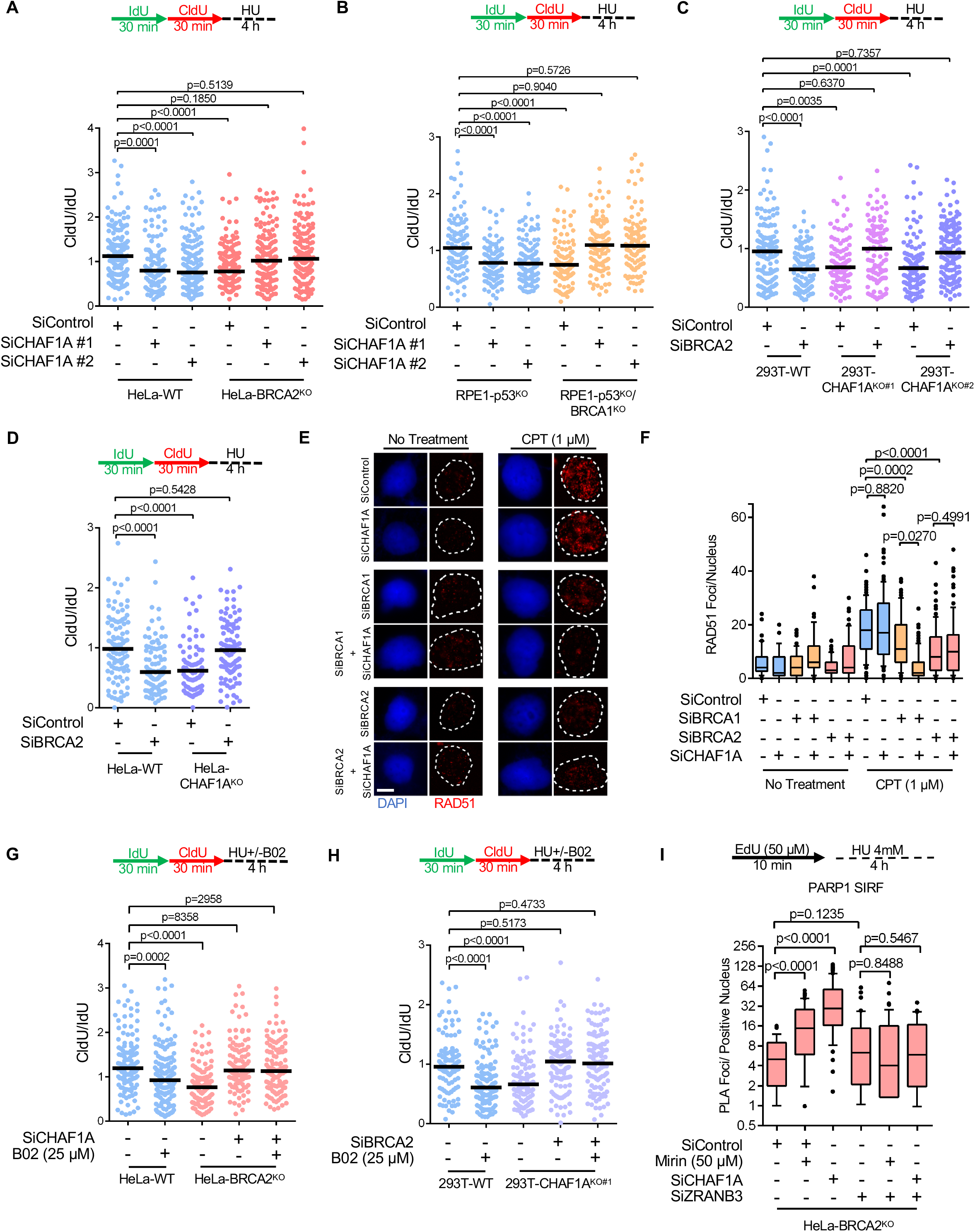
Loss of CAF-1 promotes fork stability in BRCA-deficient cells. **A, B**. DNA fiber combing assays showing that CHAF1A depletion results in HU-induced fork resection in wildtype cells, but suppresses this degradation in BRCA2-knockout HeLa cells (**A**) and in BRCA1-knockout RPE1 cells (**B**). The ratio of CldU to IdU tract lengths is presented, with the median values marked on the graph. The p-values (Mann-Whitney test) are listed at the top. Schematic representations of the DNA fiber combing assay conditions are also presented. Western blots confirming the knockdown are shown in Supplementary Fig. S1C, D. **C, D**. DNA fiber combing assays showing that CHAF1A knockout in 293T (**C**) or HeLa (**D**) cells results in HU-induced fork degradation, which is suppressed by BRCA2 knockdown. The ratio of CldU to IdU tract lengths is presented, with the median values marked on the graph. The p-values (Mann-Whitney test) are listed at the top. Schematic representations of the DNA fiber combing assay conditions are also presented. Western blots confirming the knockdown are shown in Supplementary Fig. S1G, H. **E, F**. RAD51 immunofluorescence experiment showing that CHAF1A depletion does not restore CPT-induced RAD51 foci in BRCA1 or BRCA2-depleted cells. HeLa cells were treated with 1μM CPT for 1h followed by media removal and chase in fresh media for 3h. Representative micrographs (**E**) and quantifications (**F**) are shown (scale bar represents 10μm). At least 50 cells were quantified for each condition. The median values are represented on the graph, and the p-values (Mann-Whitney test) are listed at the top. Western blots confirming the co-depletions are shown in Supplementary Fig. S1I. **G, H**. Inhibition of RAD51 by B02 treatment does not restore HU-induced fork degradation in CHAF1A-depleted HeLa-BRCA2^KO^ cells (**G**), or in BRCA2-depleted 293T-CHAF1A^KO^ cells (**H**). The ratio of CldU to IdU tract lengths is presented, with the median values marked on the graph. The p-values (Mann-Whitney test) are listed at the top. Schematic representations of the DNA fiber combing assay conditions are also presented. **I**. SIRF assay showing that PARP1 binding to nascent DNA is increased upon CHAF1A depletion in HeLa-BRCA2^KO^ cells, indicating stabilization of reversed replication forks. At least 40 positive cells were quantified for each condition. The p-values (Mann-Whitney test) are listed at the top. A schematic representation of the SIRF assay conditions is also presented. Representative micrographs (scale bar represents 10μm) are shown in Supplementary Fig. S1J. Western blots confirming the co-depletions are shown in Supplementary Fig. S1K.

Restoration of RAD51 loading on chromatin promotes fork protection in BRCA-deficient settings (Bhat et al., 2018; Clements et al., 2018). We therefore sought to assess the impact of CHAF1A inactivation on chromatin-bound RAD51 levels in BRCA1 and BRCA2-depleted cells. Upon treatment with the topoisomerase I inhibitor camptothecin (CPT), depletion of CHAF1A did not affect RAD51 foci formation in BRCA-proficient HeLa cells and failed to ameliorate the reduction in RAD51 foci formation observed in BRCA1 and BRCA2 depleted cells (Fig. 1E,F; Supplementary Fig. S1I). Treatment with the RAD51 inhibitor B02 was previously shown to elicit nascent DNA resection at stalled forks in BRCA-proficient cells (Bhat et al., 2018; Taglialatela et al., 2017). In line with this, treatment with B02 resulted in HU-induced fork degradation in wildtype HeLa and 293T cells; In contrast, B02 did not cause fork degradation in HeLa-BRCA2^KO^ cells depleted of CHAF1A (Fig. 1G), or in 293T-CHAF1A^KO^ cells depleted of BRCA2 (Fig. 1H). Collectively, these results indicate that CHAF1A inactivation restores fork stability to BRCA-deficient cells in a RAD51-independent manner.

Reversal of stalled replication forks is an essential prerequisite to nascent DNA resection in BRCA-deficient cells (Kolinjivadi et al., 2017; Lemaçon et al., 2017; Mijic et al., 2017; Taglialatela et al., 2017). Thus, we investigated if the suppression of fork degradation observed upon CHAF1A depletion in BRCA-deficient cells simply reflects a defect in fork reversal. PARP1 is a critical enabler of fork reversal (Berti et al., 2013). The presence of PARP1 at nascent DNA has been previously used as an indirect readout for fork reversal (Nieminuszczy et al., 2019). Under prolonged replication arrest, stable reversed replication forks are marked by PARP1. In contrast, forks undergoing resection lose nascent DNA on regressed arms and no longer retain the structural configuration resembling four-way junctions, thus precluding the presence of PARP1. We therefore used the SIRF (in situ detection of proteins at replication forks) assay, a proximity ligation-based approach (Roy et al., 2018; Taglialatela et al., 2017), to assess PARP1 binding to nascent DNA upon HU treatment. Similar to inhibition of MRE11 using mirin, depletion of CHAF1A increased PARP1 levels at nascent DNA in HeLa BRCA2^KO^ cells (Fig. 1I; Supplementary Fig. S1J,K). Importantly, abolishing fork reversal by depleting the fork remodeling translocase ZRANB3 restored PARP1 levels to similar levels across all conditions, indicating no baseline differences in PARP1 recruitment to nascent DNA (Fig. 1I; Supplementary Fig. S1J,K). These results indicate that loss of CHAF1A does not preclude fork reversal, but rather promotes the stability to reversed forks in BRCA-deficient cells.

### Loss of CAF-1 averts DNA damage and drives chemoresistance in BRCA-deficient cells

Previous work showed that fork degradation drives DNA damage-induced chromosomal rearrangements in BRCA-deficient cells (Ray Chaudhuri et al., 2016; Schlacher et al., 2011, 2012). Thus, we next investigated if CHAF1A inactivation could avert DNA damage accumulation in BRCA-deficient cells. We assessed replication-coupled DNA damage by immunofluorescence detection of γH2AX in cells treated with CPT, known to elicit nascent strand degradation (Taglialatela et al., 2017). Indeed, while BRCA1 and BRCA2-depleted HeLa cells exhibited increased γH2AX levels upon CPT treatment, γH2AX levels were ameliorated upon co-depletion of CHAF1A (Fig. 2A,B; Supplementary Fig. S2A). We also measured the accumulation of DNA double stranded breaks (DSBs) in these cells, using the neutral comet assay. Similar to γH2AX induction, treatment with CPT resulted in increased comet tail moments in BRCA1 and BRCA2-depleted HeLa cells, which was rescued upon co-depletion of CHAF1A (Fig. 2C,D). These findings show that restoration of fork stability to BRCA-deficient cells upon CHAF1A depletion is associated with suppression of DNA damage acumulation in these cells.

**Figure 2.**
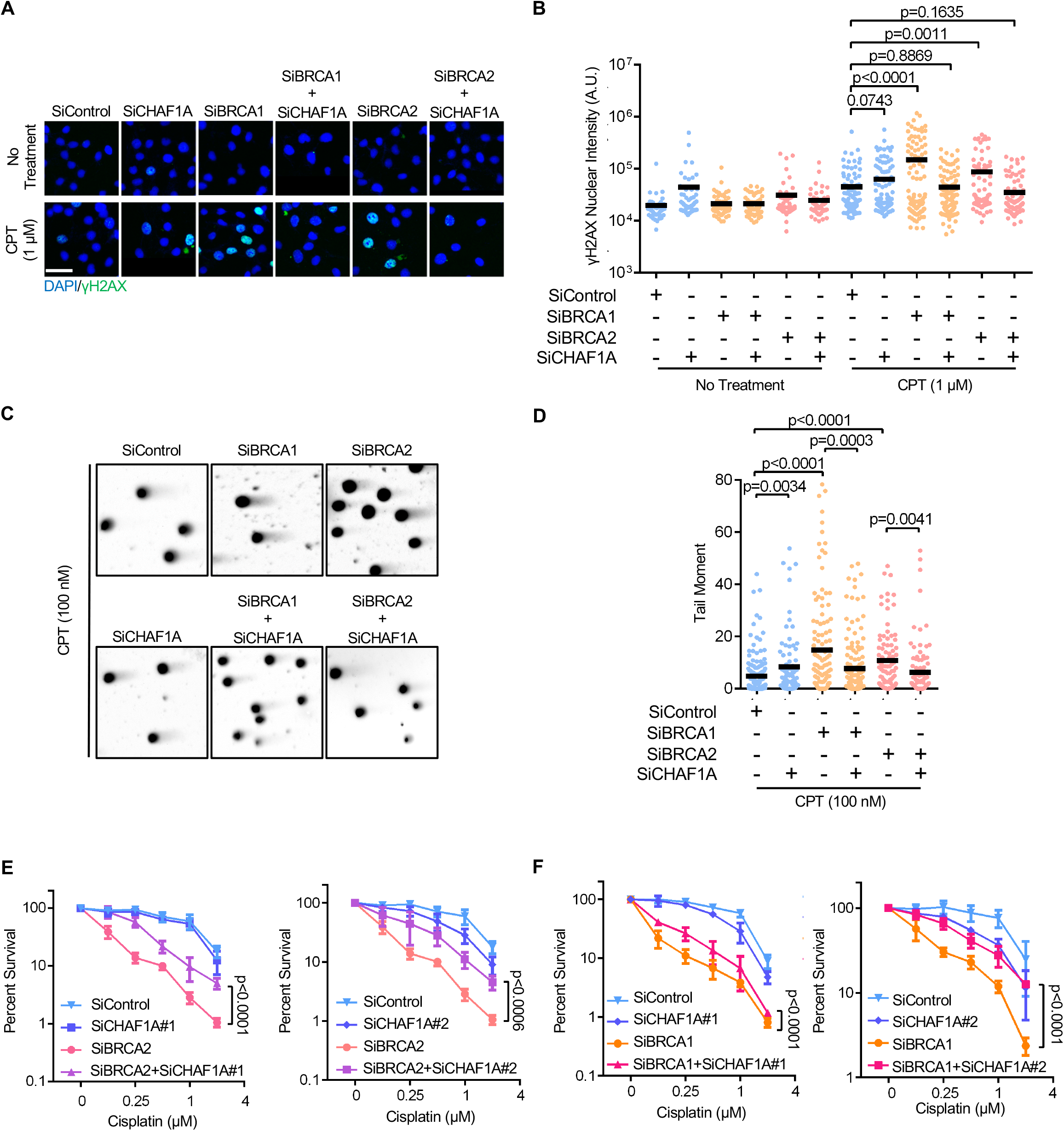
Loss of CAF-1 suppresses genomic instability in BRCA-deficient cells. **A, B**. *γ*H2AX immunofluorescence experiment showing that CHAF1A depletion suppresses CPT-induced DNA damage accumulation in BRCA1 or BRCA2-depleted cells. HeLa cells were treated with 1μM CPT for 1h followed by media removal and chase in fresh media for 3h. Representative micrographs (scale bar represents 50μm) (**A**) and quantifications (**B**) are shown. At least 50 cells were quantified for each condition. The mean values are represented on the graph, and the p-values (t-test, two-tailed, unequal variance) are listed at the top. Western blots confirming the co-depletions are shown in Supplementary Fig. S2A. **C, D**. Neutral comet assay showing that CHAF1A depletion suppresses CPT-induced DSB formation in BRCA1 or BRCA2-depleted cells. HeLa cells were treated with 100nM CPT for 4h. Representative micrographs (**C**) and quantifications (**D**) are shown. At least 85 nuclei were quantified for each condition. The mean values are represented on the graph, and the p-values (t-test, two-tailed, unequal variance) are listed at the top. **E, F**. Clonogenic survival experiments showing that CHAF1A co-depletion in BRCA2-knockdown (**E**) or BRCA1-knockdown (**F**) HeLa cells promotes cisplatin resistance. The average of three experiments, with standard deviations indicated as error bars, is shown. Asterisks indicate statistical significance (two-way ANOVA).

Restoration of fork stability is considered a driver of chemoresistance in BRCA-deficient cells (Ray Chaudhuri et al., 2016; Thakar and Moldovan, 2021). By employing clonogenic survival assays, we next investigated the impact of CHAF1A inactivation in BRCA-deficient cells on cisplatin sensitivity. CHAF1A co-depletion significantly rescued cisplatin sensitivity in Hela cells depleted of BRCA1 or BRCA2 (Fig. 2E,F). Cisplatin chemotherapy is the mainstay therapeutic approach in ovarian cancer treatment. We thus investigated if CHAF1A levels impact the chemosensitivity of BRCA-mutant ovarian tumors in clinical samples. Analyses of survival and matched genotype and expression data from TCGA datasets indicated that CHAF1A expression can stratify the survival of individuals with BRCA2-mutant ovarian tumors: high CHAF1A expression trended towards increased survival, while low CHAF1A expression trended towards reduced survival (Supplementary Fig. S2B). This is in line with our clonogenic survival results showing that CHAF1A depletion causes cisplatin resistance in BRCA2-deficient cells. Taken together, these observations suggest that CHAF1A inactivation can enable BRCA-deficient cells to avert replication-coupled DNA damage, thereby driving chemoresistance and potentially exacerbating adverse clinical outcomes in patients with BRCA1/2 mutated cancers.

### Nucleosome establishment drives replication fork protection

Since the cellular function of CAF-1 is in nucleosome deposition, we next sought to investigate if nucleosome establishment is a general determinant of fork stability in BRCA-deficient cells. The histone chaperone Anti-Silencing Factor 1 (ASF1) operates upstream of two distinct nucleosome establishment mechanisms: a CAF-1 dependent co-replicational process depositing the H3 isoform H3.1, and replication-independent processes involving the histone chaperones HIRA and DAXX, depositing the H3.3 isoform (Fromental-Ramain et al., 2017; Ransom et al., 2010; Schulz and Tyler, 2006; Tagami et al., 2004). To test if inactivating ASF1 rescues fork stability in BRCA-deficient cells, we depleted ASF1A (one of the two human ASF1 paralogs) either alone, or in conjunction with BRCA1 or BRCA2 in HeLa cells. ASF1A knockdown elicited nascent DNA resection in BRCA-proficient cells, but, in contrast to CHAF1A depletion, failed to rescue fork resection in cells depleted either of BRCA1 or BRCA2 (Fig. 3A; Supplementary Fig. S3A). This suggests that alternative ASF1-dependent nucleosome establishment pathways could compensate for CHAF1A inactivation in BRCA-deficient cells, to restore fork stability. Indeed, co-depleting CHAF1A and ASF1A in HeLa BRCA2^KO^ cells restored fork degradation, suggesting that fork stability upon CHAF1A inactivation in BRCA-deficient cells depends on ASF1 (Fig. 3B; Supplementary Fig. S3B). These results suggest that ASF1-dependent nucleosome establishment is an essential component of fork protection and determines fork stability in the context of BRCA-deficiency.

**Figure 3.**
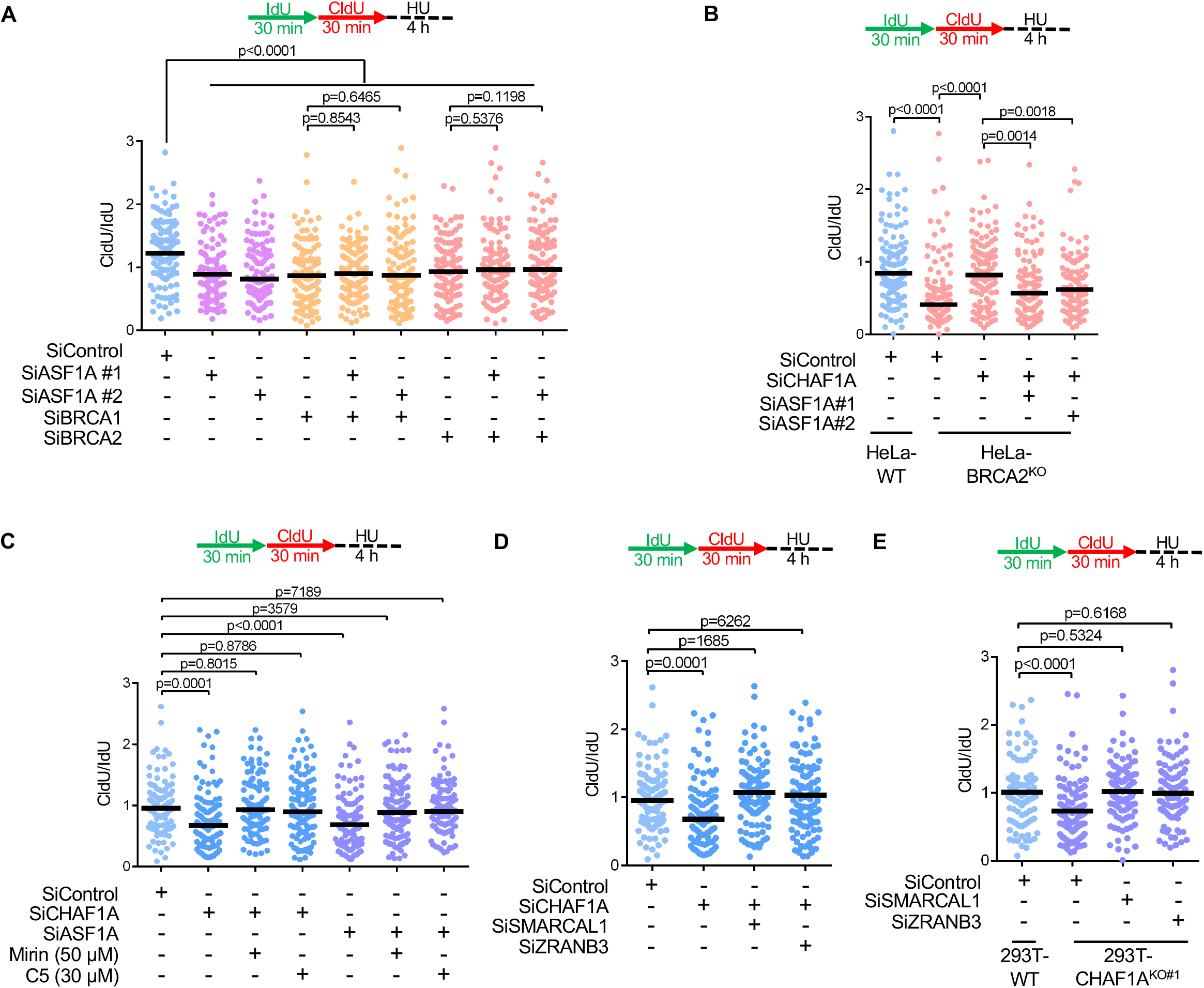
Determinants of CHAF1A-mediated fork protection. **A.** DNA fiber combing assay showing that ASF1A depletion results in HU-induced fork degradation in wildtype cells, but does not affect this degradation in BRCA1 or BRCA2-depleted HeLa cells. The ratio of CldU to IdU tract lengths is presented, with the median values marked on the graph. The p-values (Mann-Whitney test) are listed at the top. A schematic representation of the DNA fiber combing assay conditions is also presented. Western blots confirming the co-depletion are shown in Supplementary Fig. S3A. **B.** DNA fiber combing assay showing that ASF1A co-depletion restores HU-induced fork degradation in CHAF1A-knockdown HeLa-BRCA2^KO^ cells. The ratio of CldU to IdU tract lengths is presented, with the median values marked on the graph. The p-values (Mann-Whitney test) are listed at the top. A schematic representation of the DNA fiber combing assay conditions is also presented. Western blots confirming the co-depletion are shown in Supplementary Fig. S3B. **C.** DNA fiber combing assay showing that inhibition of nucleases MRE11 (by treatment with mirin) or DNA2 (by treatment with C5) suppresses HU-induced fork degradation caused by CHAF1A or ASF1A loss in HeLa cells. The ratio of CldU to IdU tract lengths is presented, with the median values marked on the graph. The p-values (Mann-Whitney test) are listed at the top. A schematic representation of the DNA fiber combing assay conditions is also presented. **D.** DNA fiber combing assay showing that co-depletion of DNA translocases SMARCAL1 and ZRANB3 suppresses HU-induced fork degradation caused by CHAF1A knockdown in HeLa cells. The p-values (Mann-Whitney test) are listed at the top. A schematic representation of the DNA fiber combing assay conditions is also presented. Western blots confirming the co-depletion are shown in Supplementary Fig. S3C. **E.** DNA fiber combing assay showing that SMARCAL1 or ZRANB3 depletion suppresses HU-induced fork degradation in 293T-CHAF1A^KO^ cells. The p-values (Mann-Whitney test) are listed at the top. A schematic representation of the DNA fiber combing assay conditions is also presented. Western blots confirming the knockdowns are shown in Supplementary Fig. S3D.

The observed epistasis between the inactivation of ASF1A and BRCA proteins led us to examine if ASF1-dependent nucleosome establishment elicits fork protection through mechanisms similar to the BRCA pathway. Loss of BRCA1 or BRCA2 function renders stalled forks susceptible to resection by the MRE11 and DNA2 nucleases (Liu et al., 2020; Ray Chaudhuri et al., 2016; Schlacher et al., 2011, 2012). Similar to this, depletion of either CHAF1A or ASF1A in HeLa cells elicited nascent DNA resection, which could be rescued by inhibition of MRE11 or DNA2 using the small molecule inhibitors mirin and C5 respectively (Fig. 3C). Next, we tested if fork reversal was also required for fork degradation in this context. Fork degradation in CHAF1A-knockdown HeLa cells, as well as in CHAF1A-knockout 293T cells, was rescued upon depletion of the fork remodeling enzymes SMARCAL1 and ZRANB3 (Fig. 3D,E; Supplementary Fig. S3C,D). These results indicate that nucleosome establishment is a general determinant of replication fork stability, and that the inactivation of nucleosome establishment elicits fork resection through mechanisms similar to those operating in BRCA-deficient cells.

### Restoration of fork stability upon CAF-1 inactivation in BRCA-deficient cells depends on the histone chaperone HIRA

ASF1A was shown to be part of two mutually-exclusive nucleosome assembly complexes involving the CAF-1 and HIRA histone chaperones, responsible for the deposition of H3.1 and H3.3 respectively (Tagami et al., 2004). HIRA-mediated histone deposition was previously shown to compensate for the inactivation of CAF-1-mediated replication-dependent nucleosome assembly (Ray-Gallet et al., 2011). Since restoration of replication fork stability is an important component of cell survival in BRCA-deficient cells (Cantor and Calvo, 2017; Thakar and Moldovan, 2021), to gain insights into a potential role for HIRA in rescuing replication fork stability, we assessed if HIRA promotes cell survival in BRCA-deficient cells upon CHAF1A inactivation. We queried publicly available CRISPR screening data for the relative dependence of BRCA1-deficient cells on HIRA and CHAF1A. We observed a linear regression of CHAF1A and HIRA gene dependency scores (CERES; lower scores correspond to higher dependencies), showing that BRCA1-proficient cells tend to dependent on both CHAF1A and HIRA for survival (Fig. 4A). This implies a general pattern of reliance on nucleosome establishment pathways. Strikingly, in cells carrying deleterious BRCA1 mutations, a lower survival dependency on CHAF1A correlated with a greater dependency on HIRA and vice-versa, suggesting that BRCA1-deficient cells rely on HIRA for cell survival in the absence of CHAF1A (Fig. 4A). It was previously shown that the histone chaperone DAXX can also cooperate with ASF1 in H3.3-dependent nucleosome assembly (Fromental-Ramain et al., 2017). In contrast to HIRA, an increased dependence on CHAF1A did not correlate with increased DAXX dependency in BRCA1-proficient cells (Fig. 4B). Moreover, in BRCA1-deficient cells, an increased dependency on DAXX was not associated with a reduced dependency on CHAF1A (Fig. 4B). These observations suggest that upon CHAF1A inactivation, BRCA-deficient cells rely on HIRA and not on DAXX to mediate nucleosome assembly and ensure cell survival.

**Figure 4.**
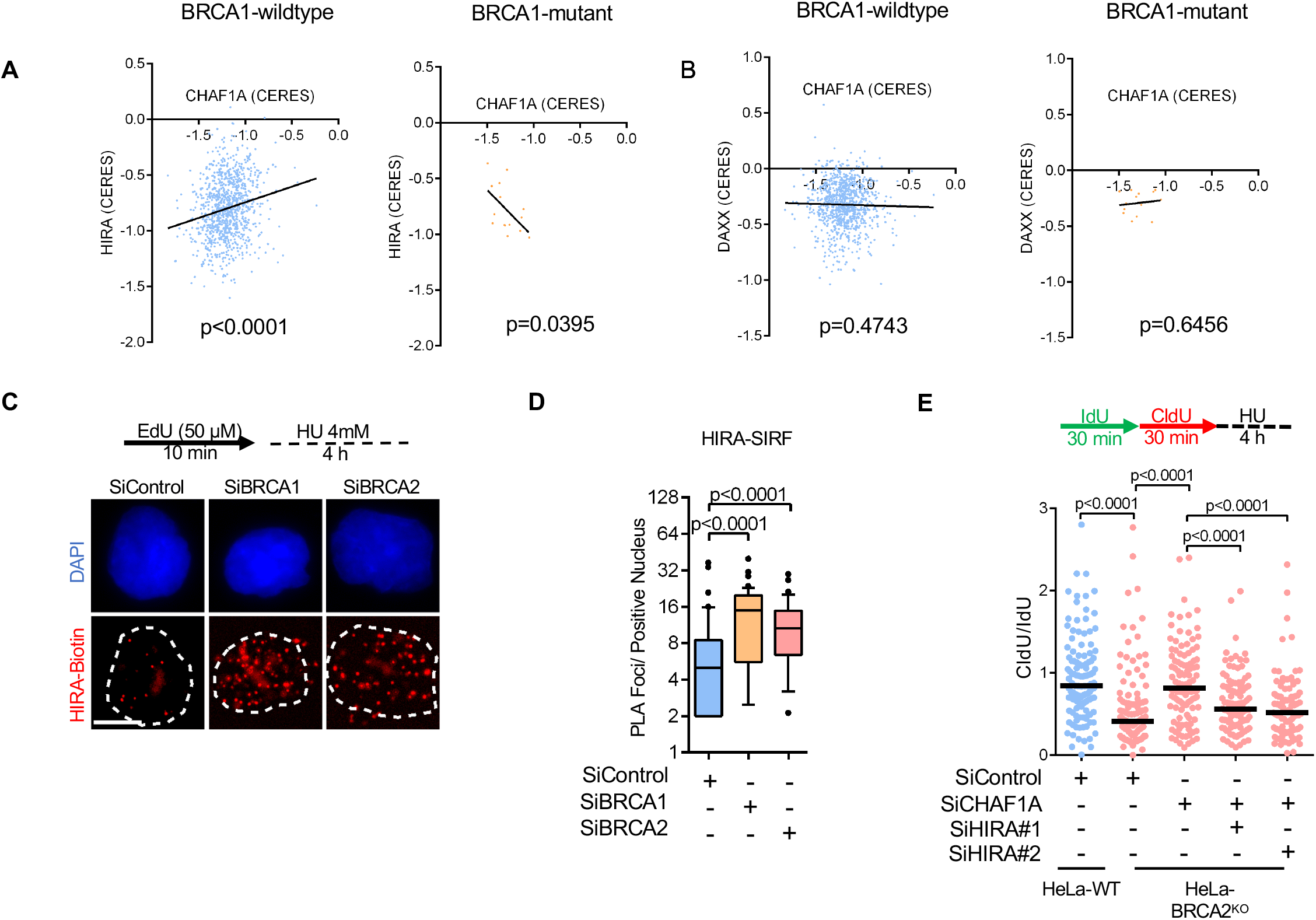
Fork protection upon CAF-1 loss in BRCA-deficient cells requires the histone chaperone HIRA. **A, B.** Linear regressions of (**A**) HIRA Gene Effect (CERES) vs. CHAF1A Gene Effect (CERES) and (**B**) DAXX Gene Effect vs. CHAF1A Gene Effect (CERES) in Cancer Cell Line Encyclopedia (CCLE) cell lines containing either wildtype BRCA1 or BRCA1 with deleterious mutations are shown. A lower CERES score corresponds to greater survival dependency. **C, D**. SIRF assay showing that HIRA binding to nascent DNA is increased upon BRCA1 or BRCA2 depletion in HeLa cells. Representative micrographs (scale bar represents 10μm) (**C**) and quantifications (**D**) are shown. At least 40 positive cells were quantified for each condition. The p-values (Mann-Whitney test) are listed at the top. A schematic representation of the SIRF assay conditions is also presented. **E.** DNA fiber combing assay showing that HIRA co-depletion restores HU-induced nascent strand resection in CHAF1A-knockdown HeLa-BRCA2^KO^ cells. The ratio of CldU to IdU tract lengths is presented, with the median values marked on the graph. The p-values (Mann-Whitney test) are listed at the top. A schematic representation of the DNA fiber combing assay conditions is also presented. Western blots confirming the co-depletion are shown in Supplementary Fig. S4D.

We next investigated if the recruitment of HIRA to stalled forks was differentially regulated in BRCA-deficient cells. SIRF assays showed that depletion of BRCA1 or BRCA2 results in increased recruitment of HIRA to HU-stalled forks (Fig. 4C,D). As a control, siRNA-mediated depletion of HIRA in HeLa cells resulted in an acute decrease in SIRF signal, confirming the specificity of this approach in detecting HIRA binding to nascent DNA (Supplementary Fig. S4A). Importantly, CHAF1A depletion had no effect on the differential recruitment of HIRA to stalled forks in BRCA-proficient and BRCA-deficient cells (Supplementary Fig. S4B), suggesting that BRCA1/2 inactivation and not CHAF1A inactivation guides the differential presence of HIRA at stalled replication forks. In contrast to HIRA, DAXX levels at stalled replication forks in BRCA1/2 depleted cells were significantly reduced (Supplementary Fig. S4C), suggesting that HIRA rather than DAXX may be operational at stalled forks in BRCA-deficient cells.

Based on these findings, we next investigated if the HIRA-H3.3 pathway of nucleosome establishment was responsible for restoring fork stability to BRCA-deficient cells upon CHAF1A loss. Indeed, co-depletion of HIRA or of H3.3 reversed the fork rescue elicited by CHAF1A knockdown in BRCA2^KO^ cells (Fig. 4E; Supplementary Fig. S4D-F). To further assess the fork-protective properties of HIRA, we depleted HIRA in wildtype HeLa cells. Unlike the depletion of ASF1A, CHAF1A, or DAXX, HIRA knockdown did not elicit nascent DNA resection upon fork stalling (Supplementary Fig. S4G,H), suggesting that HIRA-mediated fork protection is selectively activated during CHAF1A loss in BRCA-deficient cells. Altogether, these findings suggest that loss of CAF-1 triggers HIRA-mediated nucleosome establishment to protect stalled replication forks in BRCA-deficient cells.

### BRCA-deficient cells exhibit CAF-1 recycling defects during replication stress

Since HIRA-mediated fork protection in BRCA1/2-deficient cells is triggered only in the absence of CHAF1A, we sought to track the dynamics of CHAF1A at stalled forks in these cells. Replication fork stalling is accompanied by the dissociation of replisome components, including the unloading of PCNA and CAF-1 (Dungrawala et al., 2015; Yu et al., 2014). SIRF experiments showed similar CHAF1A levels at unperturbed replication forks in BRCA-proficient and BRCA1/2-depleted HeLa cells (Fig. 5A,B; Supplementary Fig. S5A). However, upon HU-induced replication arrest, BRCA-proficient cells showed lower levels of CHAF1A on EdU-labeled DNA, while CHAF1A levels remained virtually unchanged in BRCA1/2-depleted cells (Fig. 5A,B), suggesting that CAF-1 unloading from stalled forks is defective in BRCA-deficient cells.

**Figure 5.**
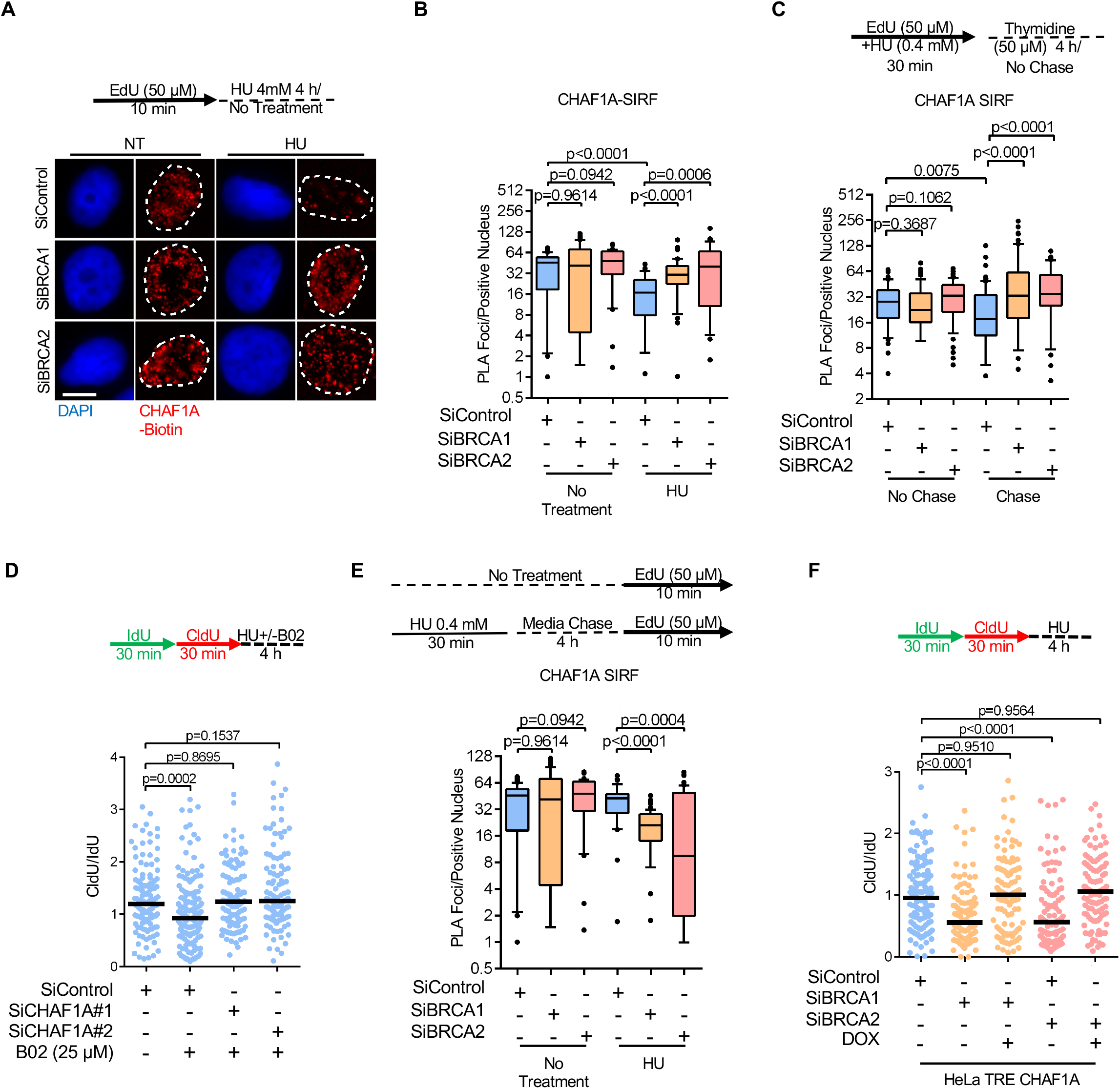
CAF-1 recycling defects underlie fork degradation in BRCA-deficient cells. **A, B.** SIRF assay showing that CHAF1A unloading from nascent DNA upon replication fork arrest is deficient in BRCA1 or BRCA2-depleted HeLa cells. To induce fork arrest, cells were treated with 4mM HU after EdU labeling. Representative micrographs (scale bar represents 10μm) (**A**) and quantifications (**B**) are shown. At least 30 positive cells were quantified for each condition. The p-values (Mann-Whitney test) are listed at the top. A schematic representation of the SIRF assay conditions is also presented. **C.** SIRF assay showing that, in BRCA1 or BRCA2-depleted HeLa cells, CHAF1A is retained on nascent DNA behind the replication fork after recovery from replication stress. Cells were labeled with EdU in the presence of low-dose HU (0.4mM for 30mins) to induce replication stress, washed, and chased for 4h in fresh media containing 50μM thymidine. At least 35 positive cells were quantified for each condition. The p-values (Mann-Whitney test) are listed at the top. A schematic representation of the SIRF assay conditions is also presented. **D.** DNA fiber combing assay showing that CHAF1A knockdown restores fork protection to wildtype HeLa cells treated with the RAD51 inhibitor B02. The ratio of CldU to IdU tract lengths is presented, with the median values marked on the graph. The p-values (Mann-Whitney test) are listed at the top. A schematic representation of the DNA fiber combing assay conditions is also presented. **E.** SIRF assay showing that prior replication stress reduces the levels of CHAF1A at ongoing replication forks in BRCA1 or BRCA2-depleted HeLa cells. Cells were subjected to low-dose HU (0.4mM for 30 mins) to induced replication stress, chased in fresh media for 4h to recover from replication stress, then labeled with EdU. At least 40 positive cells were quantified for each condition. The p-values (Mann-Whitney test) are listed at the top. A schematic representation of the SIRF assay conditions is also presented. **F.** DNA fiber combing assay showing that CHAF1A overexpression suppresses HU-induced fork degradation in BRCA1 or BRCA2-depleted HeLa cells. CHAF1A expression is under the control of the tetracycline responsive element (TRE), and is induced upon doxycycline (DOX) treatment. The ratio of CldU to IdU tract lengths is presented, with the median values marked on the graph. The p-values (Mann-Whitney test) are listed at the top. A schematic representation of the DNA fiber combing assay conditions is also presented. Western blots showing CHAF1A overexpression are presented in Supplementary Fig. S5B.

We recently showed that in PCNA ubiquitination-deficient K164R cells, the chromatin unloading of PCNA-CAF-1 complexes is defective since these complexes are retained on spontaneously accumulating ssDNA gaps on the lagging strand, which preclude Okazaki fragment maturation behind replication forks (Thakar et al., 2020). Interestingly, recent work from several laboratories has demonstrated that BRCA-deficient cells are also prone to gap accumulation during replication stress, owing to their inability to restrain replication fork progression (Cong et al., 2021; Kang et al., 2021; Liu et al., 2020; Panzarino et al., 2021). Altogether, these findings led us to hypothesize that replication stress-induced ssDNA gap accumulation in BRCA-deficient cells may retain CAF-1 on lagging strands, similar to the situation we previously described in PCNA-K164R cells. To test this, we performed SIRF experiments with EdU labeling in the presence of a low dose of HU which elicits gap formation but does not result in replication fork arrest (Panzarino et al., 2021). While the recruitment of CHAF1A to replication forks remained unchanged, BRCA1/2-depleted cells showed a persistent retention of CHAF1A at EdU-labeled DNA after a 4h thymidine chase (Fig. 5C). Importantly, CHAF1A retention defects were not observed in BRCA1/2-depleted cells under endogenous (non-HU treatment) conditions (Supplementary Fig. S5A). In contrast, the *in-situ* inactivation of RAD51 function using simultaneous treatments with B02 and HU, as opposed to the prior genetic inactivation of BRCA1/2, was enough to elicit fork rescue upon CHAF1A depletion (Fig. 5D). These findings argue that the inability of BRCA-deficient cells to restrain replication forks during replication stress gives rise to post-replicative ssDNA gaps prior to fork arrest, and drives abnormal CHAF1A retention at stalled replication forks.

We recently showed that PCNA-dependent sequestration of CHAF1A at gaps behind replication forks drives nucleosome assembly defects and fork degradation in PCNA ubiquitination-deficient K164R cells (Thakar et al., 2020). We thus hypothesized that, in BRCA-deficient cells, CAF-1 chromatin retention at replication stress-induced ssDNA gaps left behind forks, reduces its availability at ongoing replication forks; this would cause nucleosome deposition defects, and prime nascent DNA for degradation upon fork stalling. To test this, we pre-treated cells with a low dose of HU, and investigated the recruitment of CHAF1A at EdU labeled DNA after a 4h chase. CHAF1A signal was significantly lower in BRCA1/2-depleted cells than in BRCA-proficient cells (Fig. 5E), suggesting that its availability for ongoing replication forks is reduced when ssDNA gaps accumulate behind forks. Moreover, bolstering CHAF1A levels by using a doxycycline-inducible overexpression system completely rescued fork stability in cells depleted of BRCA1 and BRCA2 (Fig. 5F; Supplementary Fig. S5B). Taken together, these findings suggest that the retention of CAF-1 at ssDNA gaps behind the replication fork reduces its availability at ongoing replication forks, causing impaired nucleosome establishment which drives fork degradation in BRCA-deficient cells.

### Lagging strand gaps cause the CAF-1 recycling defects in BRCA-deficient cells

PRIMPOL-mediated repriming has recently been shown to promote ssDNA gap accumulation in BRCA-deficient cells (Kang et al., 2021; Piberger et al., 2020; Quinet et al., 2020; Simoneau et al., 2021; Taglialatela et al., 2021; Tirman et al., 2021). We therefore asked if PRIMPOL activity could potentially drive HU-induced gap accumulation in BRCA-deficient cells. We employed the BrdU alkaline comet assay to measure the accumulation of replication-associated ssDNA gaps (Mórocz et al., 2013; Thakar et al., 2020). HeLa-BRCA2^KO^ cells labeled with BrdU in the presence of a low dose of HU accumulated more replication-associated gaps compared to wildtype cells (Fig. 6A,B), in line with these recent findings. Interestingly, the depletion of PRIMPOL only partially rescued gap formation in HeLa-BRCA2^KO^ cells (Fig. 6A,B; Supplementary Fig. S6A), suggesting that a significant proportion of these gaps arise in a PRIMPOL-independent manner. Due to the continuous nature of DNA synthesis on the leading strand, it was previously suggested that PRIMPOL likely serves as a dedicated leading strand primase during replication stress (Guilliam and Doherty, 2017). We thus hypothesized that PRIMPOL-independent gaps may arise on the lagging strand in a Polα-dependent manner. Unligated Okazaki fragments result in the chromatin accumulation of poly(ADP-ribose) (PAR) chains in S-phase, which can be detected upon inhibition of the poly(ADP-ribose) glycohydrolase (PARG) enzyme (Hanzlikova et al., 2018). Indeed, SIRF experiments on cells subjected to PARG inhibition (PARGi) showed that depletion of the OF ligase LIG1 in wildtype cells results in increased PAR chain formation (Supplementary Fig. S6B,C). We reasoned that, when gaps occur on the lagging strand, the nicked DNA structure which is the substrate of LIG1 during OF ligation is not formed, since DNA synthesis on the OF is not completed. Thus, LIG1 depletion should increase PAR chain signal in cells with completed OF synthesis, but not in cells which accumulate gaps precluding nick formation. Under normal conditions, PAR SIRF experiments showed no difference in PAR chromatin levels in BRCA-proficient and BRCA1/2-depleted cells (Supplementary Fig. S6B,C). LIG1 depletion yielded a detectable increase in PAR SIRF signal, but failed to elicit differences between BRCA-proficient and BRCA1/2-depleted cells, suggesting that Okazaki fragment synthesis remains largely unperturbed in BRCA-deficient cells under normal growth conditions. We next performed SIRF to detect chromatin PAR chains after subjecting cells to EdU labeling under a low dose of HU. Interestingly, HU-induced replication stress resulted in a modest reduction in SIRF signal in BRCA1/2-depleted cells which was drastically exacerbated upon the depletion of LIG1 (Fig. 6C,D). This suggests that BRCA-deficient cells accumulate incompletely synthesized Okazaki fragments due to lagging strand gap formation upon encountering HU-induced replication stress.

**Figure 6.**
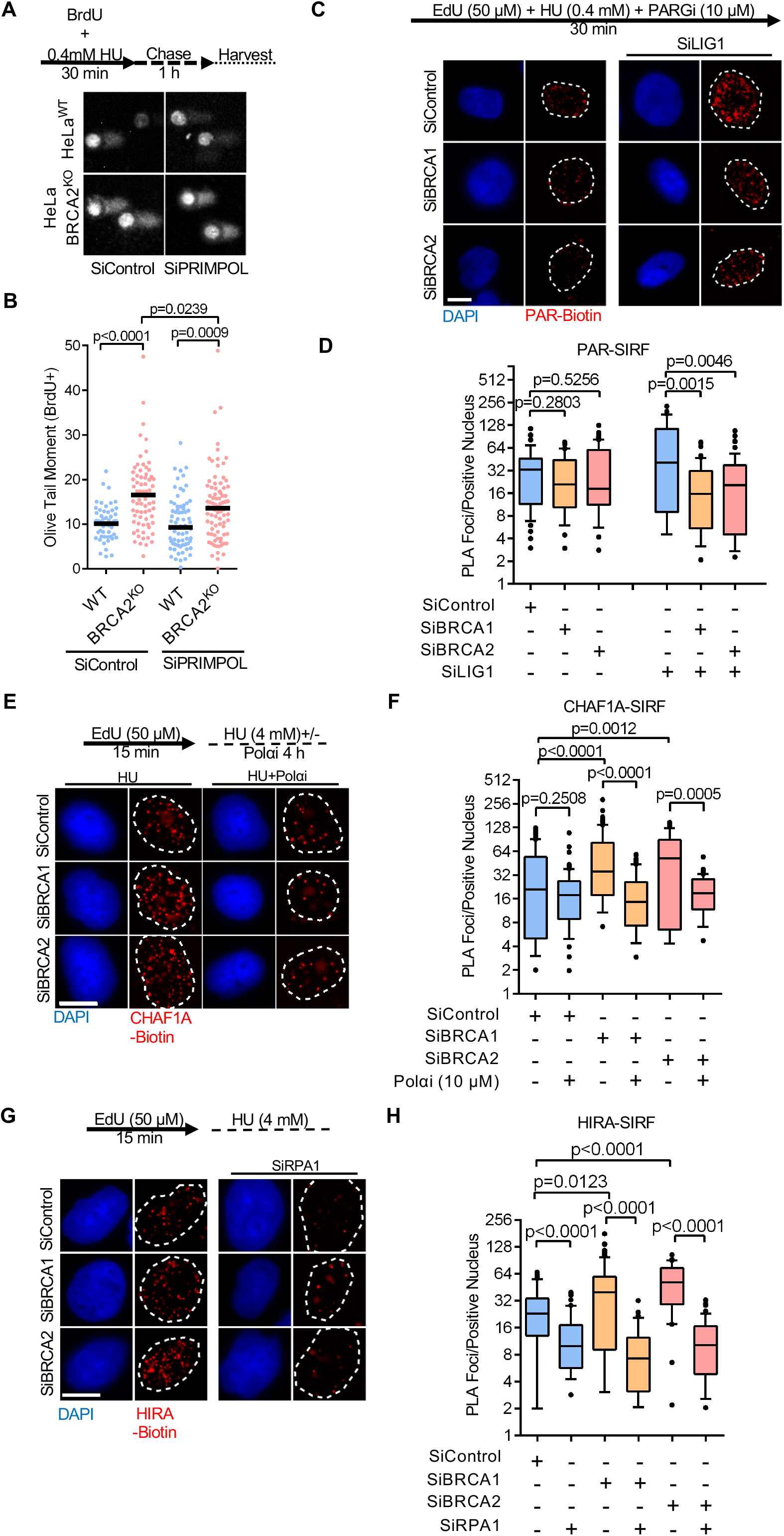
Accumulation of lagging strand gaps in BRCA-deficient cells. **A, B**. BrdU alkaline comet assay showing that PRIMPOL depletion reduces, but does not abolish replication stress-induced ssDNA accumulation in BRCA2-knockout HeLa cells, indicating the presence of PRIMPOL-independent gaps. Representative micrographs (**A**) and quantifications (**B**) are shown. At least 50 nuclei were quantified for each condition. The median values are represented on the graph, and the p-values (Mann-Whitney test) are listed at the top. A schematic representation of the assay conditions is also presented. Western blots confirming PRIMPOL depletion are presented in Supplementary Fig. S6A. **C, D.** SIRF assay showing that, upon replication stress, LIG1 knockdown induces PAR chain formation in wildtype HeLa cells, but not in BRCA1 or BRCA2-depleted HeLa cells upon treatment with a low dose of HU (0.4mM), indicating the prevalence of lagging strand gaps in these cells. Representative micrographs (scale bar represents 10μm) (**C**) and quantifications (**D**) are shown. At least 35 positive cells were quantified for each condition. The p-values (Mann-Whitney test) are listed at the top. A schematic representation of the SIRF assay conditions is also presented. Western blots confirming the co-depletion are shown in Supplementary Fig. S6C. **E, F.** SIRF assay showing that Pol*α* inhibition suppresses the increased CHAF1A retention in BRCA1 or BRCA2-depleted HeLa cells. Representative micrographs (scale bar represents 10μm) (**E**) and quantifications (**F**) are shown. At least 35 positive cells were quantified for each condition. The p-values (Mann-Whitney test) are listed at the top. A schematic representation of the SIRF assay conditions is also presented. **G, H.** SIRF assay showing that RPA1 co-depletion restores HIRA levels to the same levels in wildtype and BRCA1 or BRCA2-knockodown HeLa cells. Representative micrographs (scale bar represents 10μm) (**G**) and quantifications (**H**) are shown. At least 30 positive cells were quantified for each condition. The p-values (Mann-Whitney test) are listed at the top. A schematic representation of the SIRF assay conditions is also presented. Western blots confirming the co-depletion are shown in Supplementary Fig. S6F.

We next sought to directly test if frequent Polα-mediated repriming during transient HU-induced replication stress prior to fork stalling, could drive lagging strand gap accumulation and CAF-1 recycling defects in BRCA-deficient cells. The retinoid ST1926 was previously shown to abolish Polα activity resulting in replication fork uncoupling at sufficiently high doses (Ercilla et al., 2020). Indeed, treatment of HeLa cells with 10μM ST1926 induced maximal chromatin bound RPA levels within 5 minutes (Supplementary Fig. S6D,E), indicating a robust and immediate inhibition of Polα activity. We next performed SIRF on EdU-labeled cells subjected to a high dose of HU in the presence of ST1926. Strikingly, Polα inhibition, while having a minimal impact on BRCA-proficient cells, completely suppressed CHAF1A retention at stalled forks in BRCA1 and BRCA2-depleted cells (Fig. 6E,F). These results suggest that frequent Polα-mediated lagging strand repriming during replication stress is responsible for the CHAF1A recycling defects in BRCA-deficient cells.

Replication-associated ssDNA gaps are likely to be immediately coated by the RPA complex. Interestingly, the recruitment of HIRA to DNA during transcription has been shown to be dependent on RPA1 (Zhang et al., 2017). We wondered if RPA-coated ssDNA could account for the increased presence of HIRA observed at stalled forks in BRCA-deficient cells. Indeed, depletion of RPA1 restored HIRA at stalled forks to similar levels in both BRCA-proficient and BRCA1/2-depleted cells (Fig. 6G,H; Supplementary Fig. S6F). Taken together, these results suggest that the prevalence of lagging strand gaps, caused by Polα-dependent repriming, drives the aberrant retention of CAF-1 at stressed replication forks in BRCA-deficient cells. Furthermore, RPA complexes coating ssDNA at these gaps recruit HIRA in the proximity of HU-arrested forks in BRCA-deficient cells.

### PCNA unloading ensures CAF-1 mediated fork protection in BRCA-proficient cells

We and others previously showed that Okazaki fragment synthesis and maturation defects result in retention of PCNA-CAF-1 complexes behind replication forks (Kubota et al., 2015; Lee et al., 2013; Thakar et al., 2020). Thus, we sought to test if PCNA unloading defects cause the CAF-1 retention at stalled forks observed in BRCA-deficient cells. Indeed, similar to what we observed for CHAF1A, SIRF experiments revealed an abnormal retention of PCNA at stalled replication forks in BRCA1/2-depleted cells upon treatment with a high dose of HU (Fig. 7A,B). Importantly, Polα inhibition completely abrogated this abnormal PCNA retention, indicating that lagging strand gaps are responsible for the PCNA unloading defects in BRCA-deficient cells.

**Figure 7.**
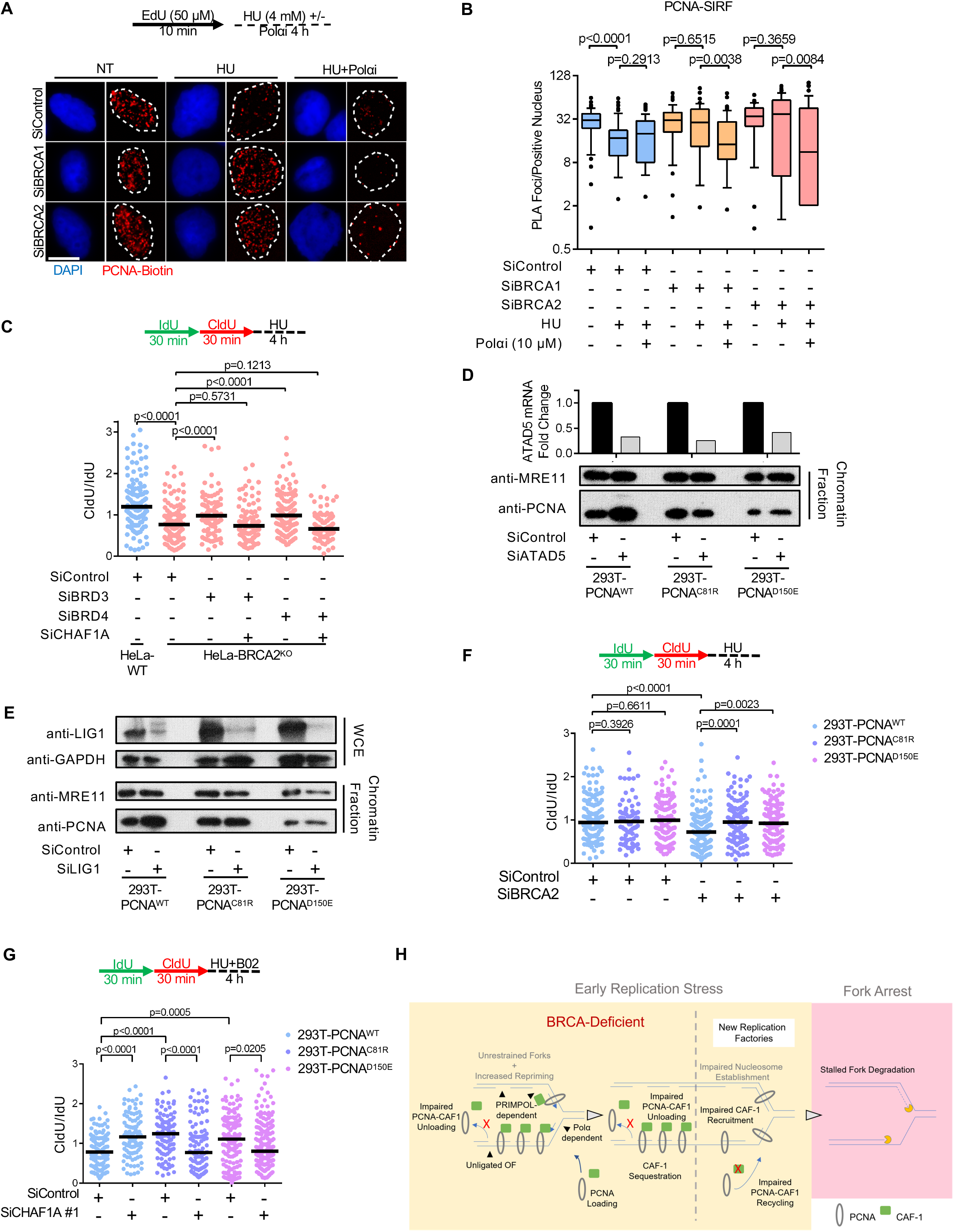
PCNA unloading defects are responsible for fork degradation in BRCA-deficient cells. **A, B.** SIRF assay showing deficient PCNA unloading from nascent DNA upon replication fork arrest in BRCA-depleted HeLa cells. Pol*α*inhibition by treatment with 10μM ST1926 suppresses this defect, indicating that this PCNA retention on nascent DNA occurs at lagging strand gaps. Representative micrographs (scale bar represents 10μm) (**A**) and quantifications (**B**) are shown. At least 50 positive cells were quantified for each condition. The p-values (Mann-Whitney test) are listed at the top. A schematic representation of the SIRF assay conditions is also presented. **C.** DNA fiber combing assay showing that co-depletion of BRD3 or BRD4 restores HU-induced fork degradation in CHAF1A-knockdown HeLa-BRCA2^KO^ cells. The ratio of CldU to IdU tract lengths is presented, with the median values marked on the graph. The p-values (Mann-Whitney test) are listed at the top. A schematic representation of the DNA fiber combing assay conditions is also presented. Western blots confirming the co-depletion are shown in Supplementary Fig. S7A. **D, E.** Chromatin fractionation experiments showing that ATAD5 (**D**) or LIG1 (**E**) knockdown increases the chromatin levels of wildtype PCNA, but not those of PCNA C81R and D150E variants. MRE11 is used as a control for the chromatin fraction. LIG1 depletion is confirmed by western blot. ATAD5 depletion is confirmed by RT-qPCR, since no antibody was available to use to verify depletion by western blot. The average of two technical replicates is shown. **F.** DNA fiber combing assay showing that BRCA2 knockdown causes HU-induced fork degradation in 293T cells expressing wildtype PCNA, but not in 293T cells expressing PCNA variants with deficient chromatin retention. The ratio of CldU to IdU tract lengths is presented, with the median values marked on the graph. The p-values (Mann-Whitney test) are listed at the top. A schematic representation of the DNA fiber combing assay conditions is also presented. Western blots confirming the knockdown are shown in Supplementary Fig. S7D. **G.** DNA fiber combing assay showing that CHAF1A knockdown suppresses HU-induced fork degradation caused by B02 treatment in 293T cells expressing wildtype PCNA, but not in 293T cells expressing PCNA variants with deficient chromatin retention. The ratio of CldU to IdU tract lengths is presented, with the median values marked on the graph. The p-values (Mann-Whitney test) are listed at the top. A schematic representation of the DNA fiber combing assay conditions is also presented. Western blots confirming the knockdown are shown in Supplementary Fig. S7E. **H.** Schematic model outlining the proposed mechanism for fork degradation in BRCA-deficient cells caused by PCNA unloading defects. The failure of BRCA-deficient cells to restrain fork progression during replication stress causes gap accumulation due to repriming by PRIMPOL (on the leading strand) and Polα (on the lagging strand). Polα-mediated repriming results in formation of lagging strand gaps which preclude PCNA unloading. The persistence of PCNA behind replication forks sequesters CAF-1 away from active replication factories, thus interfering with nucleosome assembly at forks and priming them for degradation upon stalling. A more detailed version of this model figure is provided in Supplementary Fig. S8A.

Since the results described above suggested that CAF-1 retention on ssDNA gaps promotes fork degradation in BRCA-deficient cells, we next assessed if correcting this defective PCNA unloading could restore fork stability to these cells. Recent work showed that the bromodomain extra-terminal (BET) family of proteins, namely BRD2, BRD3 and BRD4, form complexes with ATAD5 and inhibit its PCNA unloading activity (Kang et al., 2019b; Wessel et al., 2019). We reasoned that inactivating these BET proteins would enhance PCNA removal from DNA and thus reverse the effect of PCNA unloading defects in BRCA-deficient cells. Strikingly, depleting BRD3 and BRD4 restored fork stability to HeLa-BRCA2^KO^ cells (Fig. 7C; Supplementary Fig. S7A).

Importantly, co-depletion of CHAF1A with either BRD3 or BRD4 restored nascent fork degradation to HeLa-BRCA2^KO^ cells, suggesting that correction of PCNA unloading defects in BRCA-deficient cells restores fork stability in a CAF-1-dependent manner.

To more directly assess the role of PCNA unloading in ensuring fork stability in BRCA-deficient cells, we next created PCNA variants exhibiting reduced chromatin retention. We previously obtained a clonal line in pseudotriploid human 293T cells bearing genetic knockouts of four out of five endogenous PCNA alleles, with the final allele encoding a PCNA-K164R mutation (293T-PCNA^K164R(hyp)^) (Thakar et al., 2020). Importantly, we showed that these cells exhibit lower endogenous PCNA expression and the stable complementation with wildtype PCNA restored near-wildtype characteristics to these cells, thus establishing their suitability for complementation with potential PCNA mutants. Studies in *S. cerevisae* have characterized point mutations on PCNA’s trimer interface that interfere with stable homotrimer formation (Dieckman et al., 2013; Goellner et al., 2014; Kubota et al., 2015; Lau et al., 2002). We generated two of the corresponding mutations, namely PCNA-C81R and PCNA-D150E, in human PCNA (Supplementary Fig. S7B). Using a lentiviral expression system, we stably complemented 293T-PCNA^K164R(hyp)^ cells with either PCNA-WT, PCNA-C81R or PCNA-D150E variants (Supplementary Fig. S7C). We next tested if these mutations could ameliorate PCNA unloading defects caused by ATAD5 or LIG1 depletion. As expected, ATAD5 depletion increased chromatin-bound PCNA in 293T-PCNA^WT^ cells, however, 293T-PCNA^C81R^ and 293T-PCNA^D150E^ cells exhibited no increase in PCNA chromatin association upon ATAD5 depletion (Fig. 7D). Similarly, LIG1 depletion also increased chromatin-bound PCNA levels in 293T-PCNA^WT^ cells, in line with Okazaki fragment ligation being a prerequisite for PCNA unloading (Kubota et al., 2015). In contrast, LIG1 knockdown failed to cause PCNA chromatin retention in 293T-PCNA^C81R^ and 293T-PCNA^D150E^ cells (Fig. 7E). These results confirm that PCNA-C81R and PCNA-D150E mutants have reduced chromatin retention and can correct PCNA unloading defects in human cells. We next assessed if PCNA-C81R and PCNA-D150E can ameliorate fork degradation in BRCA-deficient cells. Indeed, while both BRCA2 depletion as well as RAD51 inhibition using B02 elicited fork degradation in 293T-PCNA^WT^ cells, fork stability remained intact in 293T-PCNA^C81R^ and 293T-PCNA^D150E^ cells (Fig. 7F,G; Supplementary Fig. S7D). Importantly, while CHAF1A depletion restored fork stability to B02-treated 293T-PCNA^WT^ cells, it caused fork degradation in B02-treated 293T-PCNA^C81R^ and 293T-PCNA^D150E^ cells. (Fig. 7G; Supplementary Fig. S7E). These results suggest that, in cells with impaired BRCA/RAD51 function, the restoration of PCNA unloading rescues fork stability by reinstating CAF-1 function at replication forks. Taken together, these observations imply that CAF-1 is a direct effector of the BRCA/RAD51 pathway of fork protection whose function is ensured by lagging strand gap suppression and efficient PCNA unloading (Fig. 7H; Supplementary Fig. S8A).

## DISCUSSION

### Nucleosome establishment governs BRCA-mediated fork stability

Replication-coupled nucleosome assembly is mediated by CAF-1, which operates at replication forks in a PCNA-dependent manner (Shibahara and Stillman, 1999; Zhang et al., 2000). CAF-1-mediated nucleosome assembly depends on its interaction with ASF1, which packages H3/H4 heterodimers subsequently used by CAF-1 to assemble (H3/H4)_2_ tetramers on DNA (Donham et al., 2011; Liu et al., 2012; Mattiroli et al., 2017; Sauer et al., 2017). CAF-1 preferentially interacts with the canonical histone isoform H3.1 (Tagami et al., 2004). ASF1 also participates in replication-independent deposition of histone H3.3, by cooperating with the histone chaperone HIRA (Loppin et al., 2005; Tagami et al., 2004). In humans, two ASF1 paralogs exist: ASF1A and ASF1B. Of the two, only ASF1A is capable of interacting with both CAF-1 and HIRA, and appears to do so in a mutually exclusive manner (Mello et al., 2002; Tagami et al., 2004; Tyler et al., 2001).

The presence of nucleosomes has been shown to act as a barrier to DNA resection by nucleases (Adkins et al., 2013). We previously demonstrated that CAF-1 is essential for suppression of fork degradation (Thakar et al., 2020). In the present study, we show that ASF1A inactivation is epistatic with BRCA1/2 deficiency in eliciting fork degradation (Fig. 3A). Moreover, restoring CAF-1 function at forks, through either CHAF1A overexpression or by enhancing PCNA unloading, rescued fork stability in BRCA-deficient cells (Fig. 5F; Fig. 7). Lastly, fork rescue upon CHAF1A inactivation in BRCA-deficient cells depended on the ASF1A-HIRA-H3.3 pathway of replication-independent nucleosome deposition (Fig. 3B,4E). These findings unveil nucleosome establishment as an essential effector of BRCA-mediated replication fork protection. Importantly, our findings suggest that this process is clinically relevant, since we find that CHAF1A inactivation in BRCA-deficient cells promotes chemoresistance.

Our observations indicate that the fork protective activity of HIRA only operates when CAF-1 and the BRCA pathway are simultaneously inactivated (Fig. 4E; Supplementary Fig. S4E-H). Previous studies have revealed a compensatory role of HIRA in filling nucleosome gaps on the genome in the absence of the CAF-1 (Ray-Gallet et al., 2011). However, the inactivation of CHAF1A in BRCA-proficient cells does not trigger HIRA-mediated fork protection (Fig. 1; Supplementary Fig. S1). We speculate that this selective activation of HIRA is caused by its recruitment to RPA-coated ssDNA accumulating in BRCA-deficient cells, proximal to stalled forks (Supplementary Fig. S8B). Nevertheless, the activation of HIRA-mediated fork protection still necessitates the inactivation of CHAF1A. A possible explanation for this could be the mutual exclusivity of ASF1A interactions with CAF-1 and HIRA. Indeed, ASF1 has previously been shown to ensure the supply of S-phase (H3.1-containing) histones during replication stress (Groth et al., 2005). Therefore, it is possible that the absence of CAF-1 relieves ASF1A of its S-phase histone buffering constrains, enabling it to participate in H3.3-dependent nucleosome establishment with HIRA. Furthermore, the RPA-dependent accumulation of HIRA at stalled forks in BRCA-deficient cell may prime HIRA to mediate efficient nucleosome establishment thereby preventing fork degradation upon CAF-1 loss (Supplementary Fig. S8B).

### BRCA-deficient cells accumulate Pol*α*-dependent lagging strand ssDNA gaps during replication stress

Recent studies have shown that, in BRCA-deficient cells, unrestrained replication fork progression during replication stress drives ssDNA gap accumulation through excessive PRIMPOL-dependent repriming, necessitating the engagement of post-replicative gap repair pathways, such as translesion synthesis, to avoid cellular toxicity (Cong et al., 2021; Kang et al., 2021; Liu et al., 2020; Panzarino et al., 2021; Quinet et al., 2020; Simoneau et al., 2021; Taglialatela et al., 2021; Thakar et al., 2020; Tirman et al., 2021). In these studies, the main experimental approach employed to detect replication-associated gaps involves measuring the shortening of DNA tracts upon treatment with the S1 nuclease, which specifically digests ssDNA substrates (Quinet et al., 2017b). In this assay, for a detectable shortening of replication tracts upon S1 treatment, gaps must be present on both the leading and the lagging strand. Recent work in reconstituted eukaryotic DNA replication systems revealed that fork progression is disproportionately impeded by leading strand obstacles, while lagging strand obstacles are efficiently bypassed via inherently frequent Polα mediated repriming (Guilliam and Yeeles, 2020; Taylor and Yeeles, 2018). PRIMPOL-mediated repriming is thus likely to occur mostly on the leading strand, giving rise to leading strand ssDNA gaps (Guilliam and Doherty, 2017). This raises the question of the identity of the mechanism through which lagging strand gaps occur. In this study, using an alternative method to detect ssDNA gaps, namely the BrdU alkaline comet assay, we reveal that a significant proportion of ssDNA gaps in BRCA-deficient cells accumulate independently of PRIMPOL, suggesting that they represent Polα-dependent lagging strand ssDNA gaps (Fig. 6A,B). Indeed, taking advantage of the fact that S-phase chromatin-bound PAR chains specifically form at fully synthesized but unligated OFs (Hanzlikova et al., 2018), we demonstrate that BRCA-deficient cells accumulate lagging strand discontinuities during replication stress (Fig. 6C,D). Recent evidence suggests that the BRCA pathway participates in the repair of post-replicative gaps (Cong et al., 2021; Panzarino et al., 2021; Tirman et al., 2021), suggesting that BRCA-mediated gap filling could also safeguard against lagging strand gap accumulation. Indeed, in previous work, we showed that the accumulation of ssDNA gaps in the absence of PCNA ubiquitination impairs the completion of lagging strand DNA synthesis, and necessitates BRCA-mediated gap repair (Thakar et al., 2020).

The unloading of PCNA from lagging strands can only occur upon OF ligation (Kubota et al., 2015). Importantly, we and others have shown that loss of PCNA ubiquitination, which ensures efficient lagging strand synthesis, results in PCNA unloading defects (Thakar et al., 2020; Yu et al., 2014). Thus, PCNA unloading represents a surrogate marker for OF synthesis and ligation defects. In this study, we show that PCNA unloading is defective upon replication stress in BRCA-deficient cells, but can be ameliorated by Polα inhibition. This further supports the notion that incompletely synthesized OFs accumulate in BRCA-deficient cells (Fig. 7A,B). Indeed, recent studies have also reported OF processing defects in BRCA-deficient cells (Cong et al., 2021; Paes Dias et al., 2021). In conclusion, in combination with previous studies, we provide compelling evidence that BRCA-deficient cells accumulate lagging strand gaps during replication stress.

### PCNA unloading and CAF-1 recycling connects BRCA-dependent gap suppression to fork stability

In the past decade, the inability to protect stalled forks from degradation by nucleases has emerged as a defining hallmark of BRCA-deficiency (Quinet et al., 2017a; Ray Chaudhuri et al., 2016; Schlacher et al., 2011, 2012; Thakar and Moldovan, 2021). A direct consequence of fork degradation are gross chromosomal aberrations, which likely play a crucial role in enabling tumorigenesis and cancer evolution. Furthermore, fork degradation is also associated with the hypersensitivity of BRCA-deficient cancers to chemotherapeutic agents such as cisplatin (Ray Chaudhuri et al., 2016). Conversely, the restoration of fork stability drives chemoresistance in the absence of BRCA function (Bhat et al., 2018; Ding et al., 2016; Ray Chaudhuri et al., 2016). Mechanistically, fork degradation has been shown to result in the formation of a novel substrate for endonucleolytic cleavage by MUS81, thereby enabling the restart of forks through a break-induced replication process (Lemaçon et al., 2017). However, despite having profound implications in driving genome instability in BRCA-deficient cells, fork degradation only partially explains PARPi sensitivity (Bhat et al., 2018; Cong et al., 2021; Panzarino et al., 2021; Ray Chaudhuri et al., 2016; Taglialatela et al., 2017). Instead, the inability to suppress replication-associated gaps in BRCA-deficient cells has recently emerged as a major predictor of PARPi sensitivity (Cong et al., 2021; Kang et al., 2021; Kolinjivadi et al., 2017; Nayak et al., 2020; Panzarino et al., 2021; Quinet et al., 2020; Simoneau et al., 2021; Taglialatela et al., 2021; Thakar et al., 2020). Nonetheless, how exactly ssDNA gaps underpin genome instability remains unclear. Recent evidence suggests that ssDNA gaps in BRCA-deficient cells may persist through mitosis into subsequent cell cycles, eliciting DNA damage when encountered by replication forks (Feng and Jasin, 2017; Simoneau et al., 2021; Tirman et al., 2021). In the present study, we reveal fork degradation as a direct consequence of replication-associated gap accumulation in BRCA-deficient cells. We find that the failure to restrain forks during replication stress contributes to the accumulation of lagging strand ssDNA gaps, which interferes with PCNA unloading and CAF-1 recycling in BRCA-deficient cells (Fig. 7H; Supplementary Fig. S8A). We moreover show that the reduction in CAF-1 availability at replication forks drives their degradation due to impaired replication-associated nucleosome assembly in BRCA-deficient cells.

Traditional models of BRCA-mediated fork protection assert the role of BRCA-dependent RAD51 nucleofilament assembly on reversed fork arms in acting as a direct obstacle to the action of nucleases (Kolinjivadi et al., 2017; Schlacher et al., 2011, 2012). However, recent evidence suggests that RAD51 assembly at replication forks may occur at gaps proximal to the replication forks undergoing PRIMPOL-mediated repriming (Piberger et al., 2020). Additional evidence suggests a role for the BRCA pathway in orchestrating RAD51-dependent replication fork slowing and reversal (Liu et al., 2020). Critically, fork reversal mechanisms have been shown to play an essential role in limiting PRIMPOL-mediated repriming during replication stress (Bai et al., 2020; Quinet et al., 2020; Taglialatela et al., 2021; Tirman et al., 2021). Given that we now describe a role for replication-associated gap formation in driving fork degradation, we speculate that a major mechanism by which the BRCA pathway orchestrates RAD51-mediated fork protection is by ensuring the engagement of RAD51 at ssDNA gaps proximal to, as well as behind, stressed replication forks. This enables RAD51 to suppress *de novo* ssDNA gap formation as well as rapidly repair post-replicative gaps, thereby ensuring timely PCNA unloading and CAF-1 dependent nucleosome assembly, which underlie fork protection. In conclusion, we propose a model which unifies gap suppression and fork protection, two critical functions of the BRCA pathway, and connects them to the fundamental replicative function of PCNA in orchestrating nucleosome assembly at replication forks (Fig. 7H; Supplementary Fig. S8).

## MATERIAL AND METHODS

### Cell culture and protein techniques

Human 293T, RPE1, and HeLa cells were grown in DMEM supplemented with 10% Fetal Calf Serum. For CHAF1A gene knockout, the commercially available CHAF1A CRISPR/Cas9 KO plasmid was used (Santa Cruz Biotechnology sc-402472). Transfected HeLa or 293T cells were FACS-sorted into 96-well plates using a BD FACSAria II instrument. Resulting colonies were screened by western blot. The BRCA2-knockout HeLa cells were created in our laboratory and were previously described (Clements et al., 2018). RPE1-p53^KO^-BRCA1^KO^ were obtained from Dr. Alan D’Andrea (Dana-Farber Cancer Institute, Boston, MA) (Lim et al., 2018). 293T cells with hypomorph PCNA expression were created in our laboratory and previously described (Thakar et al., 2020). For exogenous PCNA expression, pLV[Exp]-Puro-CMV lentiviral constructs encoding wildtype or the indicated variants were obtained from Cyagen. For doxycycline-induced CHAF1A overexpression, the pLV[Exp]-Puro-TRE>hCHAF1A lentiviral construct (Cyagen) was used. Infected cells were selected by puromycin.

Gene knockdown was performed using Lipofectamine RNAiMAX. AllStars Negative Control siRNA (Qiagen 1027281) was used as control. The following oligonucleotide sequences (Stealth or SilencerSelect siRNA, ThermoFisher; unless otherwise noted) were used: BRCA1: AATGAGTCCAGTTTCGTTGCCTCTG; BRCA2: GAGAGGCCTGTAAAGACCTTGAATT; SMARCAL1: CACCCTTTGCTAACCCAACTCATAA; ZRANB3: TGGCAATGTAGTCTCTGCACCTATA; ATAD5: GGTACGCTTTAAGACAGTTACTGTT; CHAF1A#1: s19499; CHAF1A#2: HSS115231; PRIMPOL: GAGGAAACCGTTGTCCTCAGTGTAT (Horizon Discovery); ASF1A#1: CAGAGAGCAGTAATCCAAATCTACA; ASF1A#2: s226043; HIRA#1: HSS111075; HIRA#2: HSS186934; DAXX: s3935; H3F3A: s51241; H3F3B: s226272; LIG1: s8173; RPA1: s12127; BRD3: s23901; BRD4: s15544.

Denatured whole cell extracts were prepared by boiling cells in 100mM Tris, 4% SDS, 0.5M β-mercaptoethanol. Chromatin fractionation was performed as previously described (Wysocka et al., 2001). Antibodies used for Western blot were: CHAF1A (Cell Signaling Technology 5480); ASF1 (Santa Cruz Biotechnology sc-53171); BRCA1 (Santa Cruz Biotechnology sc-6954); BRCA2 (Calbiochem OP95); ZRANB3 (Invitrogen PA5-65143); SMARCAL1 (Invitrogen PA5-54181); HIRA (Abcam 129169); DAXX (Invitrogen PA5-79137); RPA1 (Cell Signaling Technology 2198); LIG1 (Bethyl A301-136A); PRIMPOL (Invitrogen MA5-32899); MRE11 (GeneTex GTX70212); PCNA (Cell Signaling Technology 2586); ubiquitinated PCNA (Cell Signaling Technology 13439); BRD3 (Bethyl A302-368A); BRD4 (Bethyl A700-005); GAPDH (Santa Cruz Biotechnology sc-47724).

RAD51 and *γ*H2AX immunofluorescence was performed as we previously described (Nicolae et al., 2014). RPA2 immunofluorescence was performed as we previously described (Thakar et al., 2020). Primary antibodies used were: RAD51 (Abcam ab133534); *γ*H2AX (Millipore 05-636); RPA2 (Abcam ab2175). Secondary antibodies used were AlexaFluor 488 or AlexaFluor 568 (Invitrogen A11001, A11008, A11031, and A11036). Slides were imaged using a DeltaVision Elite confocal microscope. The number of foci/nucleus was quantified using ImageJ software.

### Drug sensitivity assays

For clonogenic survival assays, 1000 siRNA-treated cells were seeded per well in 6-well plates and incubated with the indicated doses of cisplatin. Media was changed after 1 day and colonies were stained after 10-14 days. Colonies were washed with PBS, fixed with a solution of 10% methanol and 10% acetic acid, and stained with 1% crystal violet (Aqua solutions).

Neutral and BrdU alkaline comet assays were performed as we previously described (Thakar et al., 2020), using the Comet Assay Kit (Trevigen, 4250-050) according to the manufacturer’s instructions. For the BrdU alkaline comet assay, cells were incubated with 100μM BrdU as indicated. Olive tail moment was analyzed using CometScore 2.0.

### DNA fiber assays

Cells were treated with siRNA and/or drugs according to the labeling schemes presented. Cells were incubated with 100µM IdU and 100µM CldU as indicated. Hydroxyurea and additional inhibitors (50μM mirin for MRE11 inhibition; 30μM C5 (Liu et al., 2016) for DNA2 inhibition; 25µM B02 for RAD51 inhibition; 10µM ST1926 for Polα inhibition) were added as indicated. Next, cells were collected and processed using the the FiberPrep kit (Genomic Vision EXT-001) according to the manufacturer’s instructions. DNA molecules were stretched onto coverslips (Genomic Vision COV-002-RUO) using the FiberComb Molecular Combing instrument (Genomic Vision MCS-001). Slides were then stained with antibodies detecting CldU (Abcam 6236), IdU (BD 347580), and DNA (Millipore Sigma MAD3034) and incubated with secondary Cy3, Cy5, or BV480-conjugated antibodies (Abcam 6946, Abcam 6565, and BD Biosciences 564879). Finally, the cells were mounted onto coverslips and imaged using a confocal microscope (Leica SP5). At least 70 tracts were quantified for each sample.

### Quantification of gene expression by real-time quantitative PCR (RT-qPCR)

Total mRNA was purified using TRIzol reagent (Life Tech). To generate cDNA, 1μg RNA was subjected to reverse transcription using the RevertAid Reverse Transcriptase Kit (Thermo Fisher Scientific) with oligo-dT primers. Real-time qPCR was performed with PerfeCTa SYBR Green SuperMix (Quanta), using a CFX Connect Real-Time Cycler (BioRad). The cDNA of GAPDH gene was used for normalization. Primers used were: H3F3A for: TCTGGTGCGAGAAATTGCTC; H3F3A rev: TCTTAAGCACGTTCTCCACG; H3F3B for: CGAGAGATTCGTCGTTATCAG; H3F3B rev: TGACTCTCTTAGCGTGGATG; ATAD5 for: AGGAAGAGATCCAACCAACG; ATAD5 rev: ATGTTTCGAAGGGTTGGCAG; GAPDH for: AAATCAAGTGGGGCGATGCTG; GAPDH rev: GCAGAGATGATGACCCTTTTG.

### In situ analysis of protein interactions at replication forks (SIRF)

Cells were seeded in 8-well chamber slides at 50% confluency. The following day, cells were labeled with 50µM 5-Ethynyl-2’-deoxyuridine (EdU) and treated with HU and other drugs as indicated. Cells were then extracted with 0.5% Triton-X in PBS for 10 minutes at 4°C, followed by fixation with 4% formaldehyde in PBS for 15 minutes. Fixed samples were then blocked with 3% BSA in PBS at 37°C. After blocking, samples were subjected to a Click-iT reaction with Biotin Azide for 30 minutes, followed by incubation with primary antibodies in 1% BSA and 0.05% Triton-X in PBS at 4°C overnight. Primary Antibodies used were: Biotin (mouse: Jackson ImmunoResearch 200-002-211; rabbit: Bethyl Laboratories A150-109A); PARP1 (Cell Signaling Technology 9542); CHAF1A (Cell Signaling Technology 5480); HIRA (Abcam 129169); DAXX (Invitrogen PA5-79137); PAR (R&D systems 4335-MC-100); PCNA (Cell Signaling Technology 13110). Following primary antibody treatment, cells were subjected to the PLA reaction using the Duolink kit (Millipore Sigma) according to the manufacturer’s instructions. Nuclear fluorescence signal was acquired using a DeltaVision Elite fluorescence microscope. For data analysis, cells positive for PLA fluorescence signal between biotin and the protein of interest were identified and the median foci count was represented. To control for variabilities in EdU uptake, foci counts of each sample were normalized to the respective biotin-biotin control using the geometric mean of the PLA fluorescence signal from positive cells.

### Cellular gene dependency analyses

Cellular dependency data was obtained from the DepMap Public 21Q3 dataset using the DepMap portal (depmap.org/portal). Gene knockout effects (CERES) from project Achilles CRISPR screens were obtained for BRCA1-wildtype cell lines and for cell lines bearing a deleterious mutation for BRCA1 in accordance to the Cancer Cell Line Encyclopedia Project (CCLE) (Dempster et al., 2019, 2021; Meyers et al., 2017; Pacini et al., 2021). CHAF1A and HIRA gene effects were then plotted as a regression for BRCA1-wildtype and BRCA-mutant samples. P-values for the likelihood of non-zero slopes were ascertained.

### TCGA dataset analyses

Genomic, transcriptomic, and survival data for ovarian cancer samples (Cancer Genome Atlas Research Network, 2011), part of The Cancer Genome Atlas (TCGA), were obtained from cBioPortal (Gao et al., 2013). Survival datasets were sorted by BRCA2-status and all BRCA2-mutant samples were used for subsequent analyses. Samples were divided into two groups based on CHAF1A expression status in the patient tumor samples: high (0-50^th^ percentile) and low (51^st^-100^th^ percentile). Mantel-Cox log ranked t test was used for statistical analyses of the data sets using Prism software.

### Statistical analyses

For clonogenic assays, the 2-way ANOVA statistical test was used. For immunofluorescence assays (except γH2AX staining), the DNA fiber assays, BrdU alkaline comet assays and the proximity ligation (SIRF) assays the Mann-Whitney statistical test was used. For γH2AX immunofluorescence and the neutral comet assays, the t-test (two-tailed unequal variance) was used. For immunofluorescence, DNA fiber combing, proximity ligation assays, and comet assays, results from one experiment are shown; the results were reproduced in at least one additional independent biological replicate. Statistical significance is indicated for each graph.

## ACKNOWLEDGEMENTS

We would like to thank Drs. Alan D’Andrea, Binghui Shen, Boris Pfander, and Ashna Dhoonmoon for materials and advice; and the Penn State College of Medicine Flow Cytometry and Imaging cores. This work was supported by the National Institutes of Health (R01ES026184 and R01GM134681 to GLM; R01CA244417 to CMN).

## LEGENDS TO SUPPLEMENTARY FIGURES

**Supplementary Figure S1.**
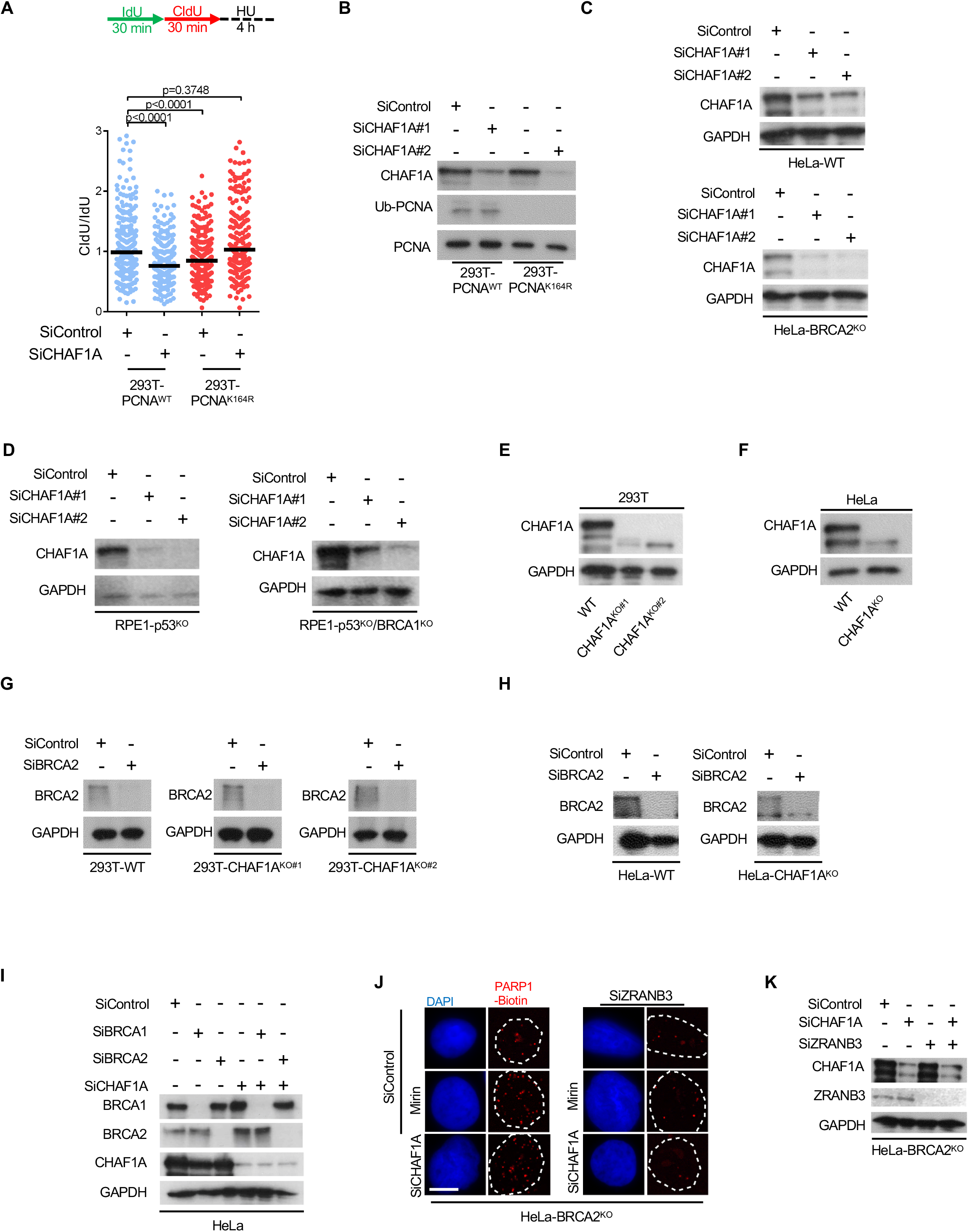
Confirmation of gene knockdowns. **A.** DNA fiber combing assays showing that CHAF1A depletion results in HU-induced fork degradation in wildtype cells, but suppresses fork degradation in 293T-PCNA^K164R^ cells. The ratio of CldU to IdU tract lengths is presented, with the median values marked on the graph. The p-values (Mann-Whitney test) are listed at the top. A schematic representation of the DNA fiber combing assay conditions is also presented. **B**. Western blots showing CHAF1A depletion in 293T cells. **C, D**. Western blots showing CHAF1A depletion in BRCA2-knockout HeLa cells (**C**) and in BRCA1-knockout RPE1 cells (**D**). **E, F**. Western blots showing CHAF1A knockout in 293T (**E**) and HeLa (**F**) cells. **G, H**. Western blots showing BRCA2 depletion in CHAF1A-knockout 293T (**G**) and HeLa (**H**) cells. **I**. Western blots showing CHAF1A co-depletion with BRCA1 or BRCA2 in HeLa cells. **J**. Representative micrographs of the PARP1 SIRF experiment (scale bar represents 10μm). **K**. Western blots showing CHAF1A co-depletion with ZRANB3 in HeLa-BRCA2^KO^ cells.

**Supplementary Figure S2.**
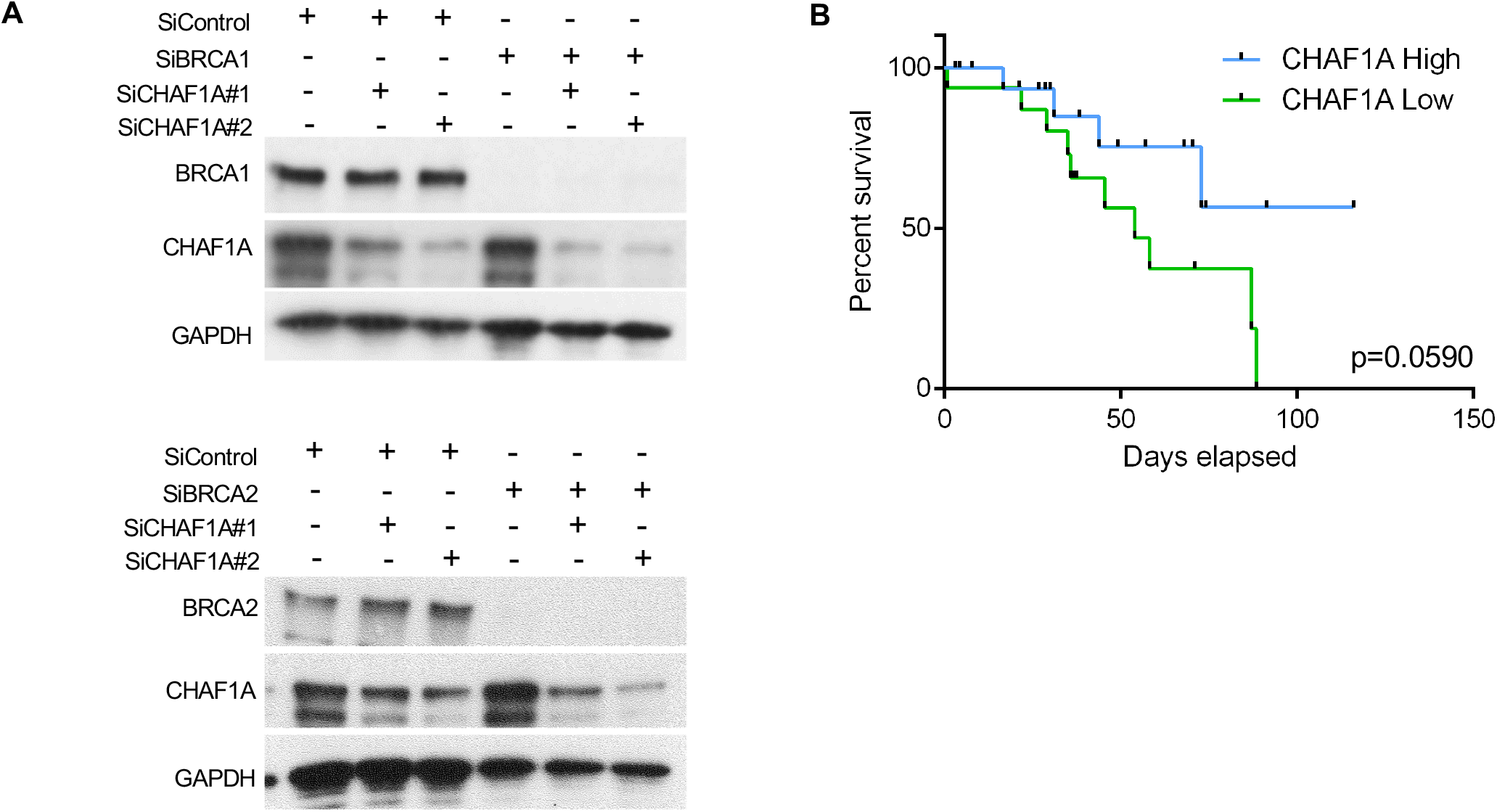
Impact of CAF-1 loss in BRCA-deficient cells. **A**. Western blots showing CHAF1A co-depletion with BRCA1 or BRCA2 in HeLa cells. **B**. Analyses of BRCA2-mutant ovarian TCGA cancer dataset showing that high CHAF1A levels are associated with increased survival, while low CHAF1A levels are associated with reduced survival. Mantel-Cox log ranked t test was used for statistical analyses (n=14, p=0.0590). The difference observed is not significant, likely because of the small number of BRCA2-mutant samples in the dataset.

**Supplementary Figure S3.**
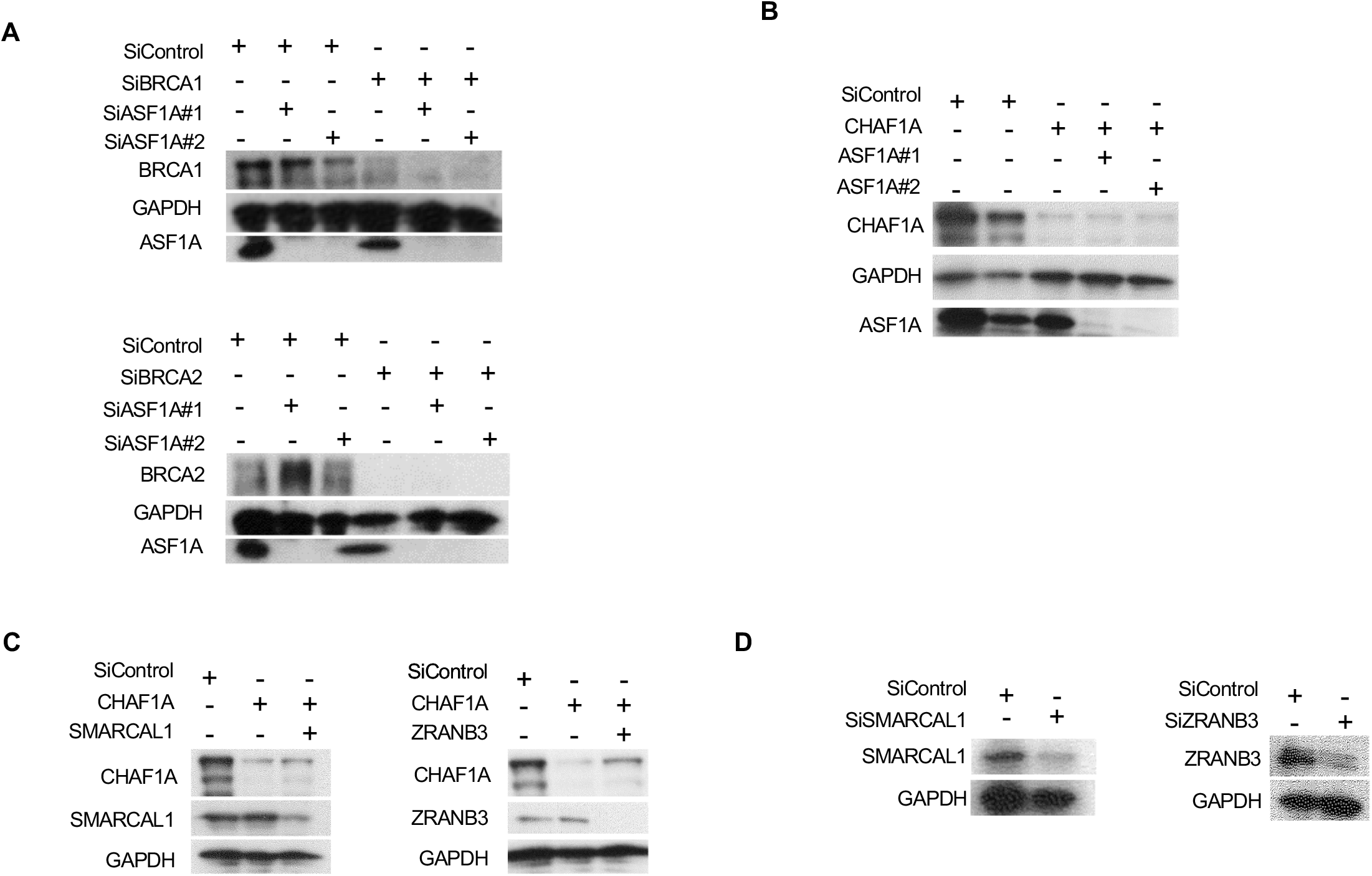
Confirmation of gene knockdowns. **A.** Western blots showing ASF1A co-depletion with BRCA1 or BRCA2 in HeLa cells. **B.** Western blots showing ASF1A co-depletion with CHAF1A in HeLa-BRCA2^KO^ cells. **C.** Western blots showing CHAF1A co-depletion with SMARCAL1 or ZRANB3 in HeLa cells. **D.** Western blots showing ZRANB3 and SMARCAL1 depletions in 293T-CHAF1A^KO^ cells.

**Supplementary Figure S4.**
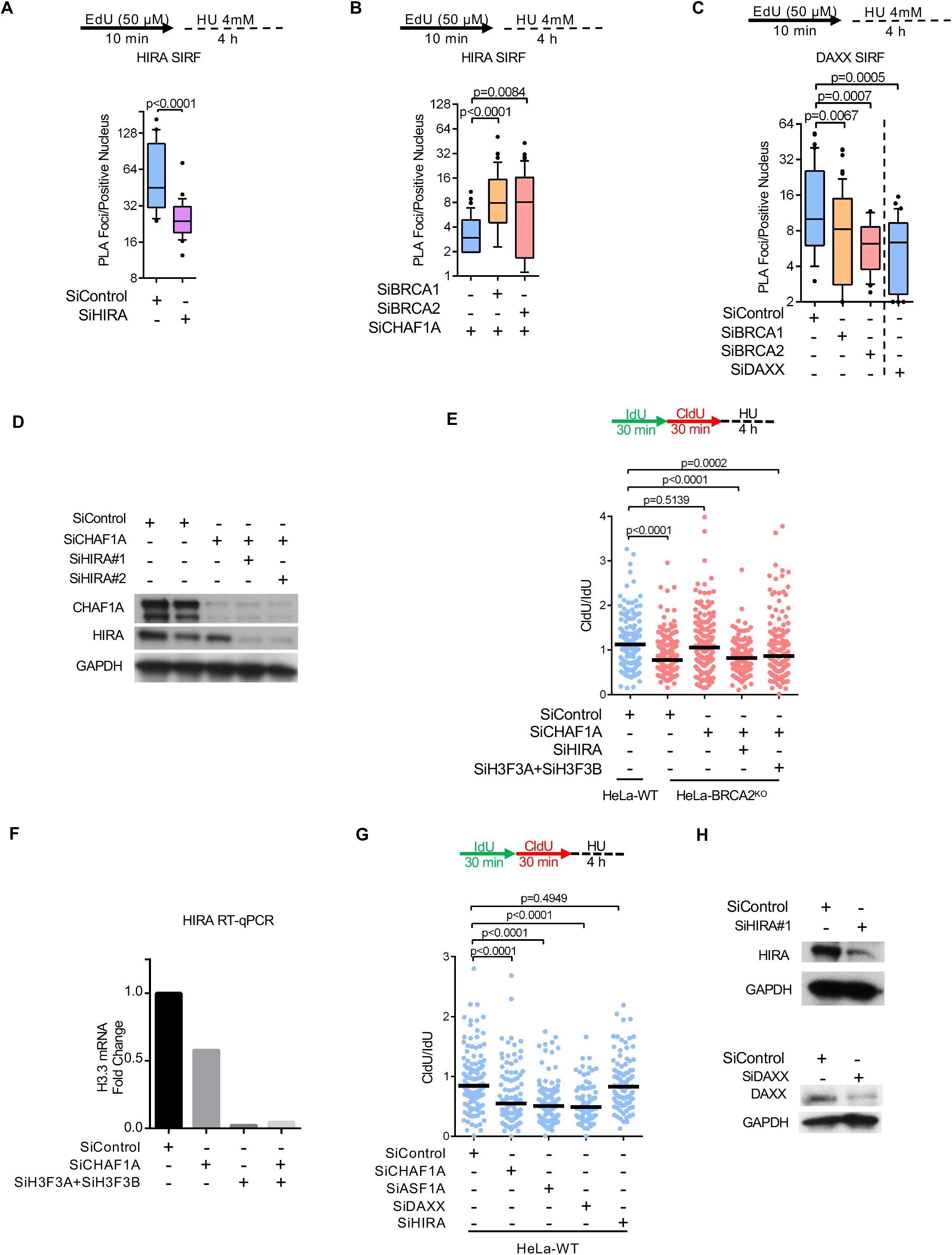
Impact of nucleosome deposition by HIRA on BRCA-deficient cells. **A**. SIRF assay confirm that the HIRA PLA signal is specific, since it is reduced upon HIRA depletion by siRNA. At least 20 positive cells were quantified for each condition. The p-values (Mann-Whitney test) are listed at the top. A schematic representation of the SIRF assay conditions is also presented. **B**. SIRF assay showing the impact of CHAF1A co-depletion on HIRA binding to nascent DNA in BRCA1 or BRCA2-knockdown HeLa cells. At least 65 cells were quantified for each condition. The p-values (Mann-Whitney test) are listed at the top. A schematic representation of the SIRF assay conditions is also presented. **C**. SIRF assay showing that DAXX binding to nascent DNA is decreased upon BRCA1 or BRCA2 depletion in HeLa cells. DAXX depletion by siRNA reduces the PLA signal, confirming its specificity. A schematic representation of SIRF assay conditions is also presented. At least 30 positive cells were quantified for each condition. The p-values (Mann-Whitney test) are listed at the top. A schematic representation of the SIRF assay conditions is also presented. **D.** Western blots showing HIRA co-depletion with CHAF1A in HeLa-BRCA2^KO^ cells. **E.** DNA fiber combing assays showing that co-depletion of HIRA or of H3.3-encoding genes H3F3A and H3F3B restores HU-induced nascent strand degradation in CHAF1A-depleted HeLa-BRCA2^KO^ cells. The ratio of CldU to IdU tract lengths is presented, with the median values marked on the graph. The p-values (Mann-Whitney test) are listed at the top. A schematic representation of the DNA fiber combing assay conditions is also presented. **F**. RT-qPCR experiment showing reduction in H3F3A and H3F3B mRNA levels upon siRNA-mediated knockdown. The average of two technical replicates is shown. (No antibody was available to us for verifying the depletion by Western blot.) **G, H.** DNA fiber combing assays showing the impact of the depletion of histone chaperones HIRA and DAXX on HU-induced nascent strand degradation in HeLa cells. The ratio of CldU to IdU tract lengths is presented, with the median values marked on the graph (**G**). The p-values (Mann-Whitney test) are listed at the top. A schematic representation of the DNA fiber combing assay conditions is also presented. Western blots confirming the knockdowns (**H**) are also presented.

**Supplementary Figure S5.**
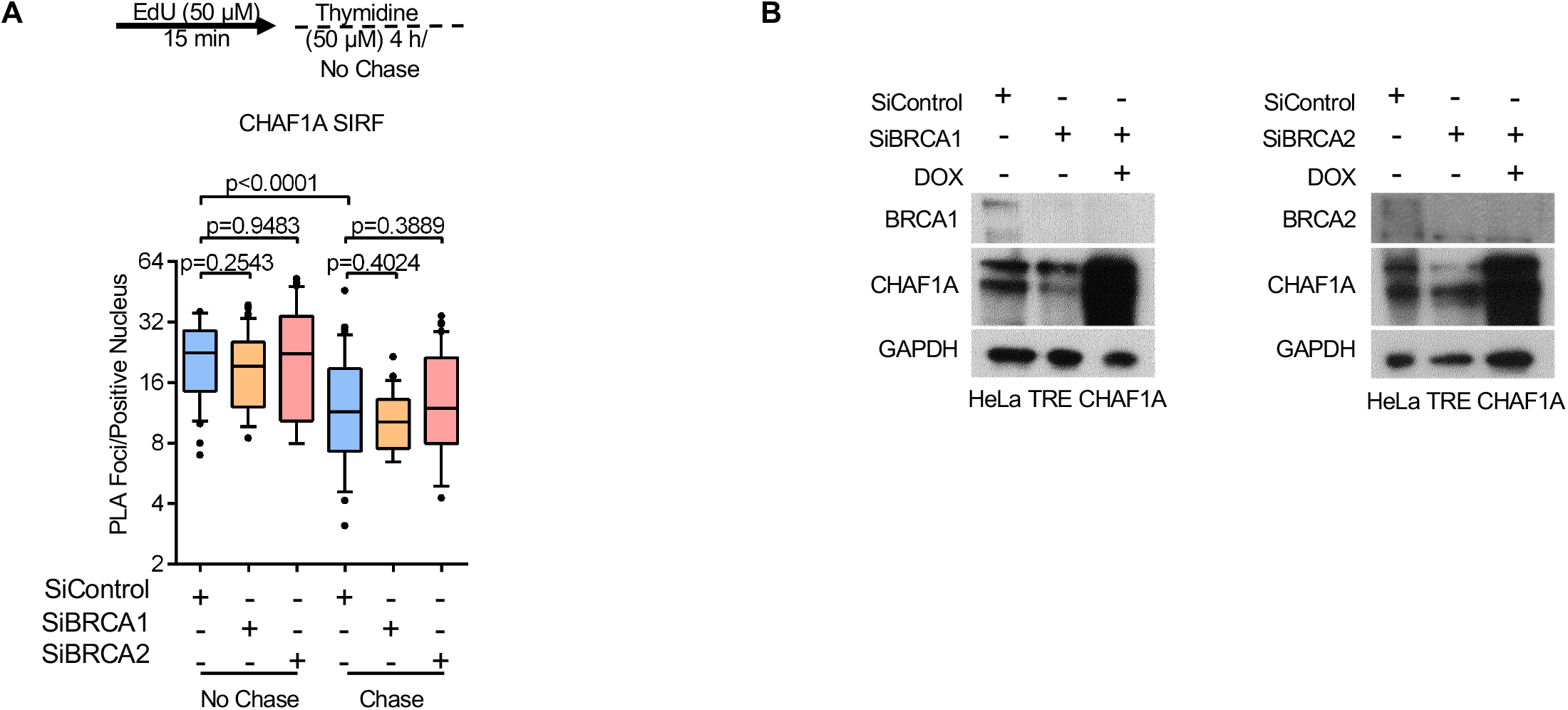
CHAF1A recycling in BRCA-deficient cells. **A.** SIRF assay showing that, in the absence of replication stress, CHAF1A recycling is not impaired in BRCA1 or BRCA2-depleted HeLa cells. Cells were labeled with EdU, washed, and chased for 4h in fresh media containing 50μM thymidine. At least 25 positive cells were quantified for each condition. The p-values (Mann-Whitney test) are listed at the top. A schematic representation of the SIRF assay conditions is also presented. **B.** Western blots showing doxycycline-induced CHAF1A overexpression, and BRCA1 or BRCA2 knockdown in these cells.

**Supplementary Figure S6.**
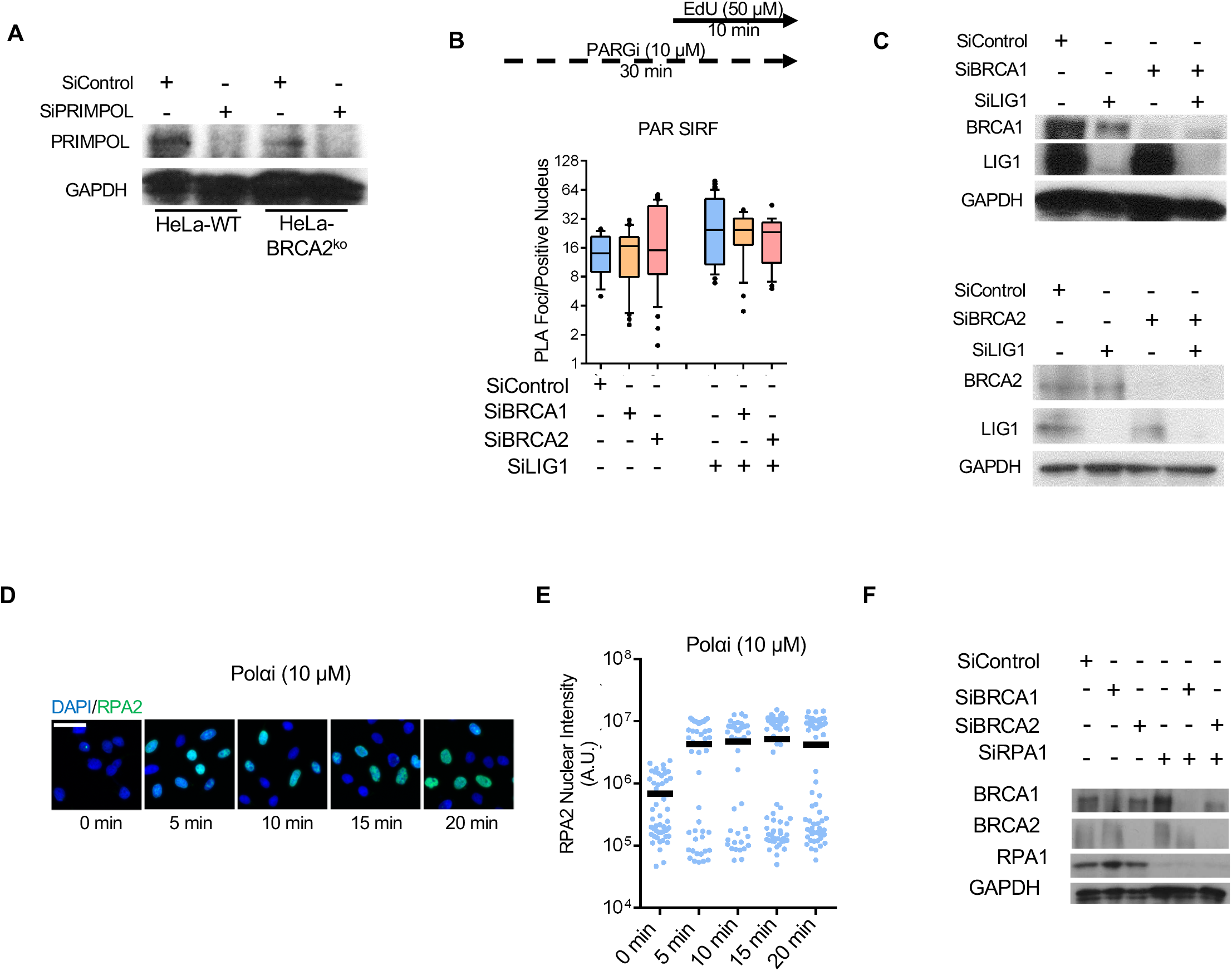
Confirmation of gene knockdowns. **A.** Western blots showing PRIMPOL depletion in HeLa wildtype or BRCA2^KO^ cells. **B.** SIRF assay showing that, under normal growth conditions (in the absence of replication stress), LIG1 knockdown induces PAR chain formation similarly in wildtype and BRCA1 or BRCA2-depleted HeLa cells. At least 25 positive cells were quantified for each condition. A schematic representation of the SIRF assay conditions is also presented. **C.** Western blots showing LIG1 co-depletion with BRCA1 or BRCA2 in HeLa cells. **D, E**. Immunofluorescence experiment showing the impact of Pol*α* inhibition by treatment with 10μM ST1926 on RPA2 chromatin foci in HeLa cells. Representative micrographs (scale bar represents 50μm) (**D**) and quantifications (**E**) are shown. At least 40 cells were quantified for each condition. The mean values are represented on the graph. **F.** Western blots showing RPA1 co-depletion with BRCA1 or BRCA2 in HeLa cells.

**Supplementary Figure S7.**
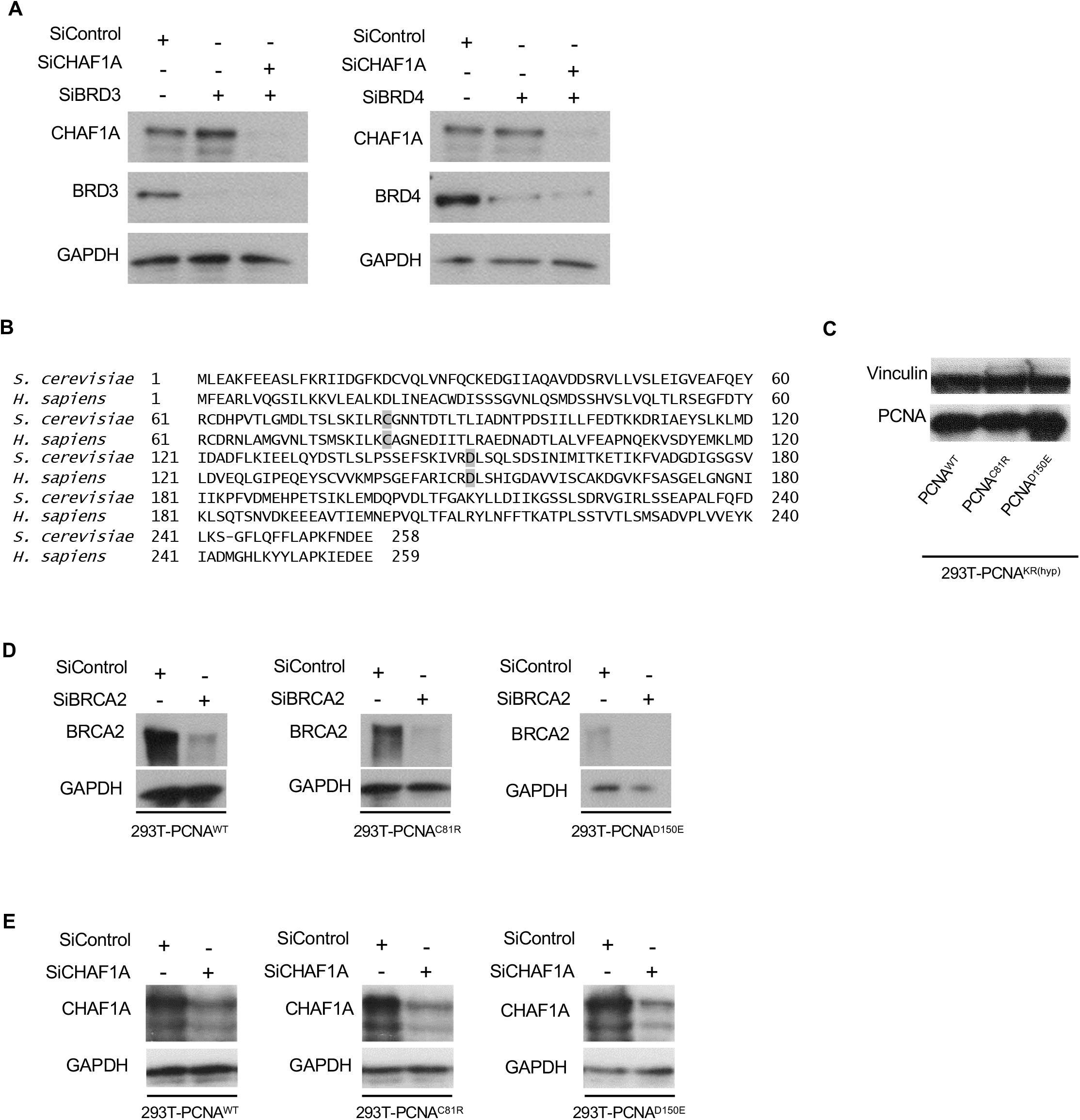
Confirmation of gene knockdowns. **A.** Western blots showing CHAF1A co-depletion with BRCA3 or BRD4 in HeLa-BRCA2^KO^ cells. **B.** Alignment of yeast and human PCNA, indicating the PCNA mutations performed. **C**. Western blots showing the expression of PCNA variants in PCNA-hypomorph 293T cells. **D.** Western blots showing BRCA2 depletion in 293T cells expressing PCNA variants. **E.** Western blots showing CHAF1A depletion in 293T cells expressing PCNA variants.

**Supplementary Figure S8.**
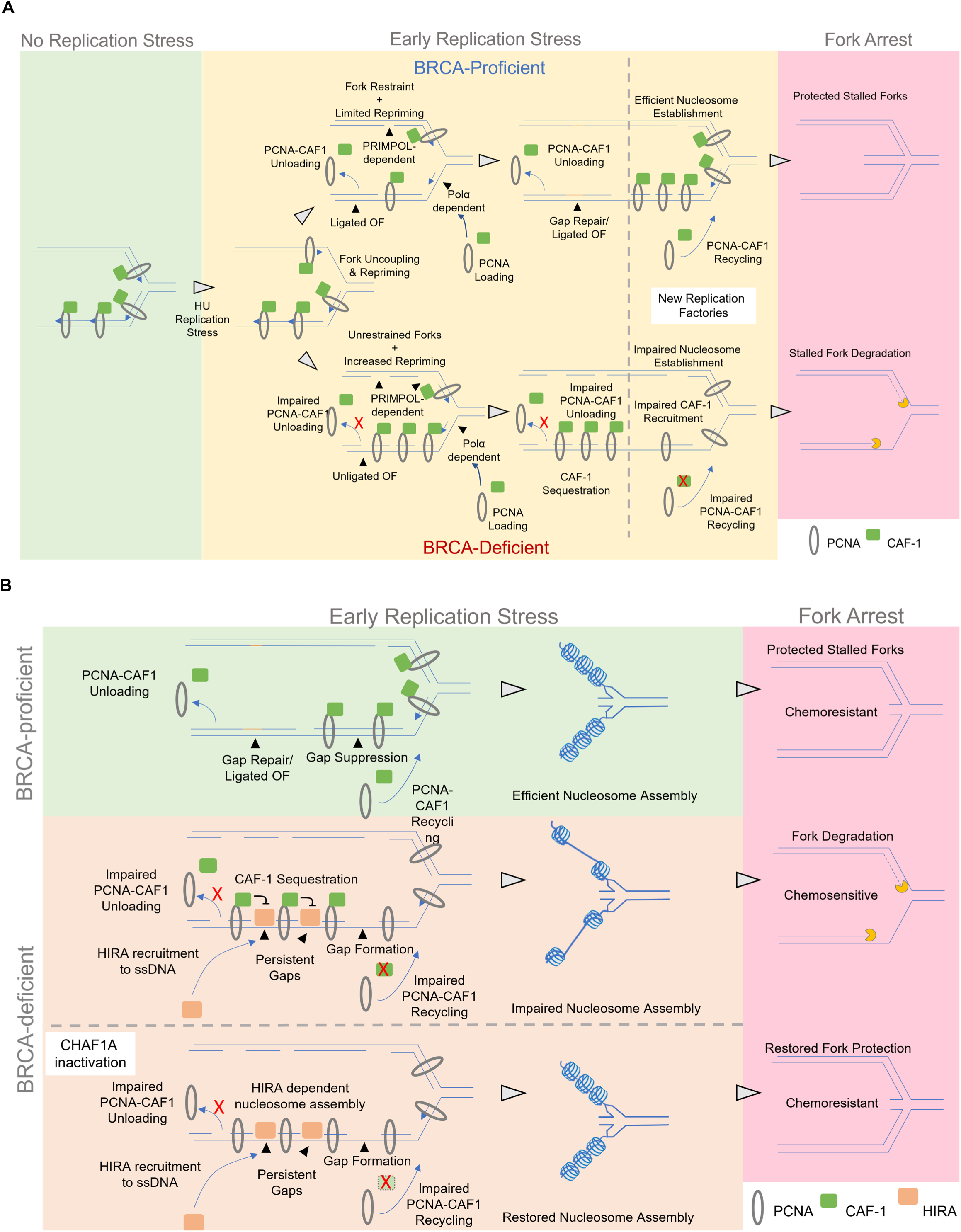
Schematic representations of the proposed models. **A.** Model for CAF-1 recycling defects as the underling factor responsible for fork degradation in BRCA-deficient cells. In BRCA-proficient cells, stressed replication forks are slowed, minimizing replication-associated gap formation. BRCA-deficient cells show impaired fork restraint during replication stress, instead undergoing repriming. On the leading strand, forks are reprimed by PRIMPOL leaving behind leading strand gaps. On the lagging strand, forks reprime through Pol*α*-mediated initiation of the subsequent OF, leaving behind lagging strand gaps. Both leading and lagging strand gaps can also be efficiently repaired by BRCA-mediated PRR. On the lagging strand, the BRCA pathway mediates gap suppression and ensures timely OF ligation, allowing unloading of PCNA-CAF-1 complexes and subsequent CAF-1 recycling to ongoing forks. This ensures proper nucleosome assembly, protecting forks from nucleolytic degradation if they arrest and reverse at a later time. In BRCA-deficient cells, gaps on the lagging strand accumulate and retain PCNA-CAF-1 complexes since OF ligation cannot take place. This reduces the availability of CAF-1 at ongoing replication forks, resulting in nucleosome deposition defects. In turn, this predisposes forks to nucleolytic degradation upon their reversal. **B.** Model for restoration of fork stability and chemoresistance upon CAF-1 loss in BRCA-deficient cells. In wildtype cells, efficient gap repair and subsequent OF ligation allows effective CAF-1 recycling and proper nucleosome assembly. In BRCA-deficient cells, ssDNA gaps accumulate and are coated by RPA, which promotes recruitment of HIRA. However, the presence of CAF-1, sequestered in PCNA complexes at lagging strand gaps, inhibits HIRA activity, possibly by competing for ASF1. Loss of CAF-1 releases HIRA activity, resulting in efficient nucleosome assembly and subsequent fork protection.

## REFERENCES

Adkins, N.L., Niu, H., Sung, P., and Peterson, C.L. (2013). Nucleosome dynamics regulates DNA processing. Nat Struct Mol Biol 20, 836–842.

Bai, G., Kermi, C., Stoy, H., Schiltz, C.J., Bacal, J., Zaino, A.M., Hadden, M.K., Eichman, B.F., Lopes, M., and Cimprich, K.A. (2020). HLTF Promotes Fork Reversal, Limiting Replication Stress Resistance and Preventing Multiple Mechanisms of Unrestrained DNA Synthesis. Mol Cell 78, 1237–1251.e7.

Balakrishnan, L., and Bambara, R.A. (2013). Okazaki Fragment Metabolism. Cold Spring Harb Perspect Biol 5, a010173.

Berti, M., Ray Chaudhuri, A., Thangavel, S., Gomathinayagam, S., Kenig, S., Vujanovic, M., Odreman, F., Glatter, T., Graziano, S., Mendoza-Maldonado, R., et al. (2013). Human RECQ1 promotes restart of replication forks reversed by DNA topoisomerase I inhibition. Nat Struct Mol Biol 20, 347–354.

Bhat, K.P., Krishnamoorthy, A., Dungrawala, H., Garcin, E.B., Modesti, M., and Cortez, D. (2018). RADX Modulates RAD51 Activity to Control Replication Fork Protection. Cell Rep 24, 538–545.

Cancer Genome Atlas Research Network (2011). Integrated genomic analyses of ovarian carcinoma. Nature 474, 609–615.

Cantor, S.B., and Calvo, J.A. (2017). Fork Protection and Therapy Resistance in Hereditary Breast Cancer. Cold Spring Harb Symp Quant Biol 82, 339–348.

Choe, K.N., and Moldovan, G.-L. (2017). Forging Ahead through Darkness: PCNA, Still the Principal Conductor at the Replication Fork. Mol Cell 65, 380–392.

Clements, K.E., Thakar, T., Nicolae, C.M., Liang, X., Wang, H.-G., and Moldovan, G.-L. (2018). Loss of E2F7 confers resistance to poly-ADP-ribose polymerase (PARP) inhibitors in BRCA2-deficient cells. Nucleic Acids Res 46, 8898–8907.

Cong, K., Peng, M., Kousholt, A.N., Lee, W.T.C., Lee, S., Nayak, S., Krais, J., VanderVere-Carozza, P.S., Pawelczak, K.S., Calvo, J., et al. (2021). Replication gaps are a key determinant of PARP inhibitor synthetic lethality with BRCA deficiency. Mol Cell 81, 3128–3144.e7.

Dempster, J.M., Rossen, J., Kazachkova, M., Pan, J., Kugener, G., Root, D.E., and Tsherniak, A. (2019). Extracting Biological Insights from the Project Achilles Genome-Scale CRISPR Screens in Cancer Cell Lines.

Dempster, J.M., Boyle, I., Vazquez, F., Root, D., Boehm, J.S., Hahn, W.C., Tsherniak, A., and McFarland, J.M. (2021). Chronos: a CRISPR cell population dynamics model.

Dieckman, L.M., Boehm, E.M., Hingorani, M.M., and Washington, M.T. (2013). Distinct structural alterations in proliferating cell nuclear antigen block DNA mismatch repair. Biochemistry 52, 5611–5619.

Ding, X., Ray Chaudhuri, A., Callen, E., Pang, Y., Biswas, K., Klarmann, K.D., Martin, B.K., Burkett, S., Cleveland, L., Stauffer, S., et al. (2016). Synthetic viability by BRCA2 and PARP1/ARTD1 deficiencies. Nat Commun 7, 12425.

Donham, D.C., Scorgie, J.K., and Churchill, M.E.A. (2011). The activity of the histone chaperone yeast Asf1 in the assembly and disassembly of histone H3/H4-DNA complexes. Nucleic Acids Res 39, 5449–5458.

Dungrawala, H., Rose, K.L., Bhat, K.P., Mohni, K.N., Glick, G.G., Couch, F.B., and Cortez, D. (2015). The replication checkpoint prevents two types of fork collapse without regulating replisome stability. Mol Cell 59, 998–1010.

Ercilla, A., Benada, J., Amitash, S., Zonderland, G., Baldi, G., Somyajit, K., Ochs, F., Costanzo, V., Lukas, J., and Toledo, L. (2020). Physiological Tolerance to ssDNA Enables Strand Uncoupling during DNA Replication. Cell Rep 30, 2416–2429.e7.

Feng, W., and Jasin, M. (2017). BRCA2 suppresses replication stress-induced mitotic and G1 abnormalities through homologous recombination. Nat Commun 8, 525.

Fromental-Ramain, C., Ramain, P., and Hamiche, A. (2017). The Drosophila DAXX-Like Protein (DLP) Cooperates with ASF1 for H3.3 Deposition and Heterochromatin Formation. Mol Cell Biol 37, e00597–16.

Gao, J., Aksoy, B.A., Dogrusoz, U., Dresdner, G., Gross, B., Sumer, S.O., Sun, Y., Jacobsen, A., Sinha, R., Larsson, E., et al. (2013). Integrative analysis of complex cancer genomics and clinical profiles using the cBioPortal. Sci Signal 6, pl1.

Goellner, E.M., Smith, C.E., Campbell, C.S., Hombauer, H., Desai, A., Putnam, C.D., and Kolodner, R.D. (2014). PCNA and Msh2-Msh6 activate an Mlh1-Pms1 endonuclease pathway required for Exo1-independent mismatch repair. Mol Cell 55, 291–304.

Groth, A., Ray-Gallet, D., Quivy, J.-P., Lukas, J., Bartek, J., and Almouzni, G. (2005). Human Asf1 regulates the flow of S phase histones during replicational stress. Mol Cell 17, 301–311.

Guilliam, T.A., and Doherty, A.J. (2017). PrimPol—Prime Time to Reprime. Genes (Basel) 8, 20.

Guilliam, T.A., and Yeeles, J.T.P. (2020). Reconstitution of translesion synthesis reveals a mechanism of eukaryotic DNA replication restart. Nat Struct Mol Biol 27, 450–460.

Hanahan, D., and Weinberg, R.A. (2011). Hallmarks of cancer: the next generation. Cell 144, 646–674.

Hanzlikova, H., Kalasova, I., Demin, A.A., Pennicott, L.E., Cihlarova, Z., and Caldecott, K.W. (2018). The Importance of Poly(ADP-Ribose) Polymerase as a Sensor of Unligated Okazaki Fragments during DNA Replication. Mol Cell 71, 319–331.e3.

Hashimoto, Y., Ray Chaudhuri, A., Lopes, M., and Costanzo, V. (2010). Rad51 protects nascent DNA from Mre11-dependent degradation and promotes continuous DNA synthesis. Nat Struct Mol Biol 17, 1305–1311.

Hoege, C., Pfander, B., Moldovan, G.-L., Pyrowolakis, G., and Jentsch, S. (2002). RAD6-dependent DNA repair is linked to modification of PCNA by ubiquitin and SUMO. Nature 419, 135–141.

Jasin, M. (2002). Homologous repair of DNA damage and tumorigenesis: the BRCA connection. Oncogene 21, 8981–8993.

Johnson, C., Gali, V.K., Takahashi, T.S., and Kubota, T. (2016). PCNA Retention on DNA into G2/M Phase Causes Genome Instability in Cells Lacking Elg1. Cell Rep 16, 684–695.

Kang, M.-S., Ryu, E., Lee, S.-W., Park, J., Ha, N.Y., Ra, J.S., Kim, Y.J., Kim, J., Abdel-Rahman, M., Park, S.H., et al. (2019a). Regulation of PCNA cycling on replicating DNA by RFC and RFC-like complexes. Nat Commun 10, 2420.

Kang, M.-S., Kim, J., Ryu, E., Ha, N.Y., Hwang, S., Kim, B.-G., Ra, J.S., Kim, Y.J., Hwang, J.M., Myung, K., et al. (2019b). PCNA Unloading Is Negatively Regulated by BET Proteins. Cell Rep 29, 4632–4645.e5.

Kang, Z., Fu, P., Alcivar, A.L., Fu, H., Redon, C., Foo, T.K., Zuo, Y., Ye, C., Baxley, R., Madireddy, A., et al. (2021). BRCA2 associates with MCM10 to suppress PRIMPOL-mediated repriming and single-stranded gap formation after DNA damage. Nat Commun 12, 5966.

Kannouche, P.L., Wing, J., and Lehmann, A.R. (2004). Interaction of human DNA polymerase eta with monoubiquitinated PCNA: a possible mechanism for the polymerase switch in response to DNA damage. Mol Cell 14, 491–500.

Karras, G.I., and Jentsch, S. (2010). The RAD6 DNA damage tolerance pathway operates uncoupled from the replication fork and is functional beyond S phase. Cell 141, 255–267.

Kolinjivadi, A.M., Sannino, V., De Antoni, A., Zadorozhny, K., Kilkenny, M., Técher, H., Baldi, G., Shen, R., Ciccia, A., Pellegrini, L., et al. (2017). Smarcal1-Mediated Fork Reversal Triggers Mre11-Dependent Degradation of Nascent DNA in the Absence of Brca2 and Stable Rad51 Nucleofilaments. Mol Cell 67, 867–881.e7.

Kubota, T., Katou, Y., Nakato, R., Shirahige, K., and Donaldson, A.D. (2015). Replication-Coupled PCNA Unloading by the Elg1 Complex Occurs Genome-wide and Requires Okazaki Fragment Ligation. Cell Rep 12, 774–787.

Kuchenbaecker, K.B., Hopper, J.L., Barnes, D.R., Phillips, K.-A., Mooij, T.M., Roos-Blom, M.-J., Jervis, S., van Leeuwen, F.E., Milne, R.L., Andrieu, N., et al. (2017). Risks of Breast, Ovarian, and Contralateral Breast Cancer for BRCA1 and BRCA2 Mutation Carriers. JAMA 317, 2402–2416.

Lau, P.J., Flores-Rozas, H., and Kolodner, R.D. (2002). Isolation and characterization of new proliferating cell nuclear antigen (POL30) mutator mutants that are defective in DNA mismatch repair. Mol Cell Biol 22, 6669–6680.

Lee, K., Fu, H., Aladjem, M.I., and Myung, K. (2013). ATAD5 regulates the lifespan of DNA replication factories by modulating PCNA level on the chromatin. J Cell Biol 200, 31–44.

Lemaçon, D., Jackson, J., Quinet, A., Brickner, J.R., Li, S., Yazinski, S., You, Z., Ira, G., Zou, L., Mosammaparast, N., et al. (2017). MRE11 and EXO1 nucleases degrade reversed forks and elicit MUS81-dependent fork rescue in BRCA2-deficient cells. Nat Commun 8, 860.

Lim, K.S., Li, H., Roberts, E.A., Gaudiano, E.F., Clairmont, C., Sambel, L.A., Ponnienselvan, K., Liu, J.C., Yang, C., Kozono, D., et al. (2018). USP1 Is Required for Replication Fork Protection in BRCA1-Deficient Tumors. Mol Cell 72, 925–941.e4.

Liu, W., Zhou, M., Li, Z., Li, H., Polaczek, P., Dai, H., Wu, Q., Liu, C., Karanja, K.K., Popuri, V., et al. (2016). A Selective Small Molecule DNA2 Inhibitor for Sensitization of Human Cancer Cells to Chemotherapy. EBioMedicine 6, 73–86.

Liu, W., Krishnamoorthy, A., Zhao, R., and Cortez, D. (2020). Two replication fork remodeling pathways generate nuclease substrates for distinct fork protection factors. Sci Adv 6, eabc3598.

Liu, W.H., Roemer, S.C., Port, A.M., and Churchill, M.E.A. (2012). CAF-1-induced oligomerization of histones H3/H4 and mutually exclusive interactions with Asf1 guide H3/H4 transitions among histone chaperones and DNA. Nucleic Acids Res 40, 11229–11239.

Lopes, M., Foiani, M., and Sogo, J.M. (2006). Multiple mechanisms control chromosome integrity after replication fork uncoupling and restart at irreparable UV lesions. Mol Cell 21, 15– 27.

Loppin, B., Bonnefoy, E., Anselme, C., Laurençon, A., Karr, T.L., and Couble, P. (2005). The histone H3.3 chaperone HIRA is essential for chromatin assembly in the male pronucleus. Nature 437, 1386–1390.

Majka, J., and Burgers, P.M.J. (2004). The PCNA-RFC families of DNA clamps and clamp loaders. Prog Nucleic Acid Res Mol Biol 78, 227–260.

Mattiroli, F., Gu, Y., Yadav, T., Balsbaugh, J.L., Harris, M.R., Findlay, E.S., Liu, Y., Radebaugh, C.A., Stargell, L.A., Ahn, N.G., et al. (2017). DNA-mediated association of two histone-bound complexes of yeast Chromatin Assembly Factor-1 (CAF-1) drives tetrasome assembly in the wake of DNA replication. Elife 6, e22799.

Mello, J.A., Silljé, H.H.W., Roche, D.M.J., Kirschner, D.B., Nigg, E.A., and Almouzni, G. (2002). Human Asf1 and CAF-1 interact and synergize in a repair-coupled nucleosome assembly pathway. EMBO Rep 3, 329–334.

Meyers, R.M., Bryan, J.G., McFarland, J.M., Weir, B.A., Sizemore, A.E., Xu, H., Dharia, N.V., Montgomery, P.G., Cowley, G.S., Pantel, S., et al. (2017). Computational correction of copy number effect improves specificity of CRISPR-Cas9 essentiality screens in cancer cells. Nat Genet 49, 1779–1784.

Mijic, S., Zellweger, R., Chappidi, N., Berti, M., Jacobs, K., Mutreja, K., Ursich, S., Ray Chaudhuri, A., Nussenzweig, A., Janscak, P., et al. (2017). Replication fork reversal triggers fork degradation in BRCA2-defective cells. Nat Commun 8, 859.

Mórocz, M., Gali, H., Raskó, I., Downes, C.S., and Haracska, L. (2013). Single Cell Analysis of Human RAD18-Dependent DNA Post-Replication Repair by Alkaline Bromodeoxyuridine Comet Assay. PLoS One 8, e70391.

Nayak, S., Calvo, J.A., Cong, K., Peng, M., Berthiaume, E., Jackson, J., Zaino, A.M., Vindigni, A., Hadden, M.K., and Cantor, S.B. (2020). Inhibition of the translesion synthesis polymerase REV1 exploits replication gaps as a cancer vulnerability. Sci Adv 6, eaaz7808.

Neelsen, K.J., and Lopes, M. (2015). Replication fork reversal in eukaryotes: from dead end to dynamic response. Nat Rev Mol Cell Biol 16, 207–220.

Nicolae, C.M., Aho, E.R., Vlahos, A.H.S., Choe, K.N., De, S., Karras, G.I., and Moldovan, G.-L. (2014). The ADP-ribosyltransferase PARP10/ARTD10 interacts with proliferating cell nuclear antigen (PCNA) and is required for DNA damage tolerance. J Biol Chem 289, 13627–13637.

Nieminuszczy, J., Broderick, R., Bellani, M.A., Smethurst, E., Schwab, R.A., Cherdyntseva, V., Evmorfopoulou, T., Lin, Y.-L., Minczuk, M., Pasero, P., et al. (2019). EXD2 Protects Stressed Replication Forks and Is Required for Cell Viability in the Absence of BRCA1/2. Mol Cell 75, 605–619.e6.

Pacini, C., Dempster, J.M., Boyle, I., Gonçalves, E., Najgebauer, H., Karakoc, E., van der Meer, D., Barthorpe, A., Lightfoot, H., Jaaks, P., et al. (2021). Integrated cross-study datasets of genetic dependencies in cancer. Nat Commun 12, 1661.

Paes Dias, M., Tripathi, V., van der Heijden, I., Cong, K., Manolika, E.-M., Bhin, J., Gogola, E., Galanos, P., Annunziato, S., Lieftink, C., et al. (2021). Loss of nuclear DNA ligase III reverts PARP inhibitor resistance in BRCA1/53BP1 double-deficient cells by exposing ssDNA gaps. Mol Cell S1097-2765(21)00739-5.

Panzarino, N.J., Krais, J.J., Cong, K., Peng, M., Mosqueda, M., Nayak, S.U., Bond, S.M., Calvo, J.A., Doshi, M.B., Bere, M., et al. (2021). Replication Gaps Underlie BRCA Deficiency and Therapy Response. Cancer Res 81, 1388–1397.

Piberger, A.L., Bowry, A., Kelly, R.D.W., Walker, A.K., González-Acosta, D., Bailey, L.J., Doherty, A.J., Méndez, J., Morris, J.R., Bryant, H.E., et al. (2020). PrimPol-dependent single-stranded gap formation mediates homologous recombination at bulky DNA adducts. Nat Commun 11, 5863.

Quinet, A., Lemaçon, D., and Vindigni, A. (2017a). Replication Fork Reversal: Players and Guardians. Mol Cell 68, 830–833.

Quinet, A., Carvajal-Maldonado, D., Lemacon, D., and Vindigni, A. (2017b). DNA Fiber Analysis: Mind the Gap! Methods Enzymol 591, 55–82.

Quinet, A., Tirman, S., Jackson, J., Šviković, S., Lemaçon, D., Carvajal-Maldonado, D., González-Acosta, D., Vessoni, A.T., Cybulla, E., Wood, M., et al. (2020). PRIMPOL-Mediated Adaptive Response Suppresses Replication Fork Reversal in BRCA-Deficient Cells. Mol Cell 77, 461–474.e9.

Ransom, M., Dennehey, B.K., and Tyler, J.K. (2010). Chaperoning histones during DNA replication and repair. Cell 140, 183–195.

Ray Chaudhuri, A., Callen, E., Ding, X., Gogola, E., Duarte, A.A., Lee, J.-E., Wong, N., Lafarga, V., Calvo, J.A., Panzarino, N.J., et al. (2016). Replication fork stability confers chemoresistance in BRCA-deficient cells. Nature 535, 382–387.

Ray-Gallet, D., Woolfe, A., Vassias, I., Pellentz, C., Lacoste, N., Puri, A., Schultz, D.C., Pchelintsev, N.A., Adams, P.D., Jansen, L.E.T., et al. (2011). Dynamics of histone H3 deposition in vivo reveal a nucleosome gap-filling mechanism for H3.3 to maintain chromatin integrity. Mol Cell 44, 928–941.

Roy, S., Luzwick, J.W., and Schlacher, K. (2018). SIRF: Quantitative in situ analysis of protein interactions at DNA replication forks. J Cell Biol 217, 1521–1536.

Sauer, P.V., Timm, J., Liu, D., Sitbon, D., Boeri-Erba, E., Velours, C., Mücke, N., Langowski, J., Ochsenbein, F., Almouzni, G., et al. (2017). Insights into the molecular architecture and histone H3-H4 deposition mechanism of yeast Chromatin assembly factor 1. Elife 6, e23474.

Schlacher, K., Christ, N., Siaud, N., Egashira, A., Wu, H., and Jasin, M. (2011). Double-strand break repair-independent role for BRCA2 in blocking stalled replication fork degradation by MRE11. Cell 145, 529–542.

Schlacher, K., Wu, H., and Jasin, M. (2012). A distinct replication fork protection pathway connects Fanconi anemia tumor suppressors to RAD51-BRCA1/2. Cancer Cell 22, 106–116.

Schulz, L.L., and Tyler, J.K. (2006). The histone chaperone ASF1 localizes to active DNA replication forks to mediate efficient DNA replication. FASEB J 20, 488–490.

Scully, R., and Livingston, D.M. (2000). In search of the tumour-suppressor functions of BRCA1 and BRCA2. Nature 408, 429–432.

Shibahara, K., and Stillman, B. (1999). Replication-dependent marking of DNA by PCNA facilitates CAF-1-coupled inheritance of chromatin. Cell 96, 575–585.

Simoneau, A., Xiong, R., and Zou, L. (2021). The trans cell cycle effects of PARP inhibitors underlie their selectivity toward BRCA1/2-deficient cells. Genes Dev 35, 1271–1289.

Stelter, P., and Ulrich, H.D. (2003). Control of spontaneous and damage-induced mutagenesis by SUMO and ubiquitin conjugation. Nature 425, 188–191.

Tagami, H., Ray-Gallet, D., Almouzni, G., and Nakatani, Y. (2004). Histone H3.1 and H3.3 complexes mediate nucleosome assembly pathways dependent or independent of DNA synthesis. Cell 116, 51–61.

Taglialatela, A., Alvarez, S., Leuzzi, G., Sannino, V., Ranjha, L., Huang, J.-W., Madubata, C., Anand, R., Levy, B., Rabadan, R., et al. (2017). Restoration of Replication Fork Stability in BRCA1- and BRCA2-Deficient Cells by Inactivation of SNF2-Family Fork Remodelers. Mol Cell 68, 414–430.e8.

Taglialatela, A., Leuzzi, G., Sannino, V., Cuella-Martin, R., Huang, J.-W., Wu-Baer, F., Baer, R., Costanzo, V., and Ciccia, A. (2021). REV1-Polζ maintains the viability of homologous recombination-deficient cancer cells through mutagenic repair of PRIMPOL-dependent ssDNA gaps. Mol Cell 81, 4008–4025.e7.

Taylor, M.R.G., and Yeeles, J.T.P. (2018). The Initial Response of a Eukaryotic Replisome to DNA Damage. Mol Cell 70, 1067–1080.e12.

Thakar, T., and Moldovan, G.-L. (2021). The emerging determinants of replication fork stability. Nucleic Acids Res 49, 7224–7238.

Thakar, T., Leung, W., Nicolae, C.M., Clements, K.E., Shen, B., Bielinsky, A.-K., and Moldovan, G.-L. (2020). Ubiquitinated-PCNA protects replication forks from DNA2-mediated degradation by regulating Okazaki fragment maturation and chromatin assembly. Nat Commun 11, 2147.

Tirman, S., Quinet, A., Wood, M., Meroni, A., Cybulla, E., Jackson, J., Pegoraro, S., Simoneau, A., Zou, L., and Vindigni, A. (2021). Temporally distinct post-replicative repair mechanisms fill PRIMPOL-dependent ssDNA gaps in human cells. Mol Cell 81, 4026–4040.e8.

Tyler, J.K., Collins, K.A., Prasad-Sinha, J., Amiott, E., Bulger, M., Harte, P.J., Kobayashi, R., and Kadonaga, J.T. (2001). Interaction between the Drosophila CAF-1 and ASF1 chromatin assembly factors. Mol Cell Biol 21, 6574–6584.

Welcsh, P.L., Owens, K.N., and King, M.C. (2000). Insights into the functions of BRCA1 and BRCA2. Trends Genet 16, 69–74.

Wessel, S.R., Mohni, K.N., Luzwick, J.W., Dungrawala, H., and Cortez, D. (2019). Functional Analysis of the Replication Fork Proteome Identifies BET Proteins as PCNA Regulators. Cell Rep 28, 3497–3509.e4.

Wysocka, J., Reilly, P.T., and Herr, W. (2001). Loss of HCF-1-chromatin association precedes temperature-induced growth arrest of tsBN67 cells. Mol Cell Biol 21, 3820–3829.

Yu, C., Gan, H., Han, J., Zhou, Z.-X., Jia, S., Chabes, A., Farrugia, G., Ordog, T., and Zhang, Z. (2014). Strand-specific analysis shows protein binding at replication forks and PCNA unloading from lagging strands when forks stall. Mol Cell 56, 551–563.

Zellweger, R., Dalcher, D., Mutreja, K., Berti, M., Schmid, J.A., Herrador, R., Vindigni, A., and Lopes, M. (2015). Rad51-mediated replication fork reversal is a global response to genotoxic treatments in human cells. J Cell Biol 208, 563–579.

Zhang, H., Gan, H., Wang, Z., Lee, J.-H., Zhou, H., Ordog, T., Wold, M.S., Ljungman, M., and Zhang, Z. (2017). RPA Interacts with HIRA and Regulates H3.3 Deposition at Gene Regulatory Elements in Mammalian Cells. Mol Cell 65, 272–284.

Zhang, Z., Shibahara, K., and Stillman, B. (2000). PCNA connects DNA replication to epigenetic inheritance in yeast. Nature 408, 221–225.

